# Functional characterisation of an essential neo-chromosome III in Sc2.0 strain reveals opportunities and challenges for genome minimisation in Sc3.0

**DOI:** 10.64898/2026.04.20.719597

**Authors:** Reem Swidah, Marco Monti

## Abstract

Large-scale genome minimisation in eukaryotes remains a major challenge due to essential genes embedded within deletion-refractory regions and pervasive synthetic lethal interactions. Here, we address these limitations by engineering an essential neo–chromosome III that relocates all 14 essential genes from synthetic chromosome III onto a separate chromosome, thereby enabling further minimisation of synIII. To further expand design space, we created highly synthetic neo-chromosome variants with sequences absent from natural genomes. We refactored essential gene expression using both native and orthogonal promoter–terminator pairs from *Saccharomyces paradoxus* and *S. eubayanus*. Reporter assays showed that orthogonal regulatory elements largely recapitulate *S. cerevisiae* activity. Both architectures restored viability in essential gene deletion libraries, demonstrating robust cross-species complementation. Engineered linear and circular forms of essential neo-chromosomes were highly stable over 100 generations and supported a near wild-type phenotype. Relocating essential functions enabled SCRaMbLE-mediated deletion of previously inaccessible regions, substantially expanding the deletion landscape. To improve the screening efficiency of SCRaMbLEd strains, we developed a SCRaMbLE reporter, ERICA (Elementary Random Integration Cassette), a loxPsym-flanked *URA3* cassette that integrates randomly and enables iterative selection. Nanopore sequencing confirmed complex rearrangements, including deletions of up to ∼40 kb and loss of essential loci. Together, this work establishes a modular and extensible platform for orthogonal essential gene engineering and SCRaMbLE-enabled genome reduction, providing key design principles for next-generation synthetic eukaryotic genomes. These findings have broad implications beyond yeast, providing transferable design principles for genome minimisation in more complex eukaryotic systems, including mammalian and human cells.

## Introduction

The genomes of living organisms have evolved over approximately 3.8 billion years, resulting in complex structures across multiple levels of organisation. Despite major advances in molecular biology, this complexity is still not fully understood. This challenge arises for two main reasons. First, genomes have evolved to support survival and reproduction under diverse environmental pressures. However, many genes appear non-essential under laboratory conditions, and the functions of some genes remain unknown (Kobayashi et al., 2003; Peña-Castillo & Hughes, 2007; Serres et al., 2001). *[1–3].* For example, in *Saccharomyces cerevisiae*, over 1000 of the 6000 genes, have unknown functions, serve as redundancy, and are condition-specific [2–4]. Secondly, gene regulation and genetic interactions are highly complex, even in well-studied microorganisms such as *S. cerevisiae* [5]. Scientists have begun integrating biology with cutting-edge engineering techniques not only to understand biology better but also to leverage engineering biology to address global challenges such as pharmaceutical manufacturing, biofuel production, and environmental conservation. However, the complexity of biological systems limits our ability to modify existing genomes, potentially leading to instability of heterologous pathways, genetic incompatibility, and low product yields due to competition for cellular resources [6]. With significant advancements in DNA synthesis, synthesizing entire microbial genomes has become achievable and cost-effective. For instance, in 2002, the poliovirus cDNA (∼7.5 kb) was the first to be chemically synthesized [7]. In 2003, the ϕX174 bacteriophage genome was synthesized without using actual DNA as a template [8]. In 2008, Gibson et al. synthesized *Mycoplasma genitalium*, the first prokaryotic smallest bacterial genome with a 582 kb genome, but it has a very slow growth rate [9]. Following that, a faster-growing strain of *M. mycoides* (JCVI-syn1.0) was completed in 2010 [10]. In 2019, the Jason W. Chin group engineered a recoded bacterial genome by removing 3 codons, resulting in a synthetic *E. coli* that uses 61 codons. This strain has the potential to produce proteins with novel functions through codon reassignment [11]. The most altered synthetic genome to date is the synthetic genome of the Sc2.0 yeast strain of *S. cerervisiae.* The Sc2.0 project aims to create a fully synthetic genome of *S. cerevisiae*, which is nearing completion. A 6.5 synthetic yeast strain was successfully generated in J. Boekés lab in 2023 [12–18]. In this project, multiple design features are introduced to the genome such as: removing all the elements that cause genome instability such as introns and transposon while all tRNA genes are relocated to a tRNA Neo-chromosome [19]; incorporating PCRTag within the ORF as a watermark allowing synthetic genes to be differentiated from the WT genes counterpart; replacing all TAG stop codon to TAA which paves the way to study genetic code expansion. Importantly, one of the key features of the Sc2.0 project is the insertion of symmetrical loxP sites downstream of every non-essential gene. Upon Cre recombinase induction ***S****ynthetic **C**hromosome **R**earrangement **a**nd **M**odification **b**y **L**oxP-mediated **E**volution* (SCRaMbLE) takes place. SCRaMbLE shuffles the synthetic genome and introduces combinatorial rearrangements including deletions, duplications, inversions, and insertions, evolving the synthetic yeast strain toward a desired phenotype through the creation of large genotype libraries [13, 15, 17]. Reducing genome complexity by constructing a minimal genome has become feasible and holds the potential to unveil the secrets of biology at the genome level. Genome minimization aims to create a genome with a minimal set of genes necessary for survival/life under defined conditions. compared to its wild-type counterpart. A minimal genome strain, with fewer uncharacterized elements and a reduced complex regulatory network, is expected to be easier to study and engineer simultaneously. Engineering a minimal genome would enhance our understanding and unlock the fundamentals of genome biology. Insights derived from the synthetic minimal genome could identify genomic regions that could be removed, defragmented, and refactored, enabling the building of epigenetic regulation through genome redesign. Moreover, simplified microorganisms harboring minimal genomes would serve as robust chassis for diverse biotechnological applications. They would enable a better understanding and engineering of biological systems by reducing genome complexity. This reduction improves the stability of genetic devices and heterologous metabolic pathways by minimising unwanted genomic interactions. It also enhances predictability by simplifying regulatory and metabolic networks, facilitating more accurate modelling. Eliminating accessory genes, proteins, and pathways can free up biosynthetic capacity. Ultimately, such systems would allow us to address fundamental biological questions and create new-to-nature genomes that were previously unattainable. [6]. The challenge in defining a minimal genome lies in creating it in the first place. A synthetic minimal genome is defined as a combination of essential genes and a minimal gene set that sustains life under specific conditions [6]. Essential genes, which carry out essential biological functions, are the genes required for the survival of a cell or organism [20]. The budding yeast *S. cerevisiae* contains approximately 6,000 open reading frames, ∼20% of which are essential. Essential genes can be classified into four categories. Firstly, conditional essential genes are required for survival only under specific environmental or nutritional conditions; for example, the auxotrophic marker *URA3* is conditionally essential, as its deletion causes lethality unless uracil is supplemented in the growth medium [21]. Secondly, essential genes are required for cell survival under optimal growth conditions; for instance, *ACS2*, encoding acetyl-CoA synthetase, is essential for fermentative growth on glucose, and null mutants are inviable in rich media [22, 23]. Thirdly, redundant essential genes (synthetic lethal/sick) are genes whose individual loss is tolerated due to functional redundancy, but whose combined deletion results in lethality or severe growth defects; an example is the paralogous genes *BDF1* and *BDF2*, encoding bromodomain-containing transcription factors, where single deletions are viable but double deletion is lethal [24, 25]. A single deletion does not affect cell viability, but double deletions cause lethality [26]. Finally, absolute essential genes (minimal genes) are those necessary and sufficient to sustain cellular life under optimal, stress-free conditions; these include genes involved in DNA replication, repair, protein translation, and central metabolism [27, 28]. One of the key focuses of synthetic biology is to simplify the chassis and create synthetic minimal cells, which serve as platforms to integrate functional synthetic parts, devices, and systems with functions that cannot be found in nature (Zhang et al., 2010). In prokaryotes, there are two approaches to minimizing the genome: a bottom-up approach based on chemical synthesis and a top-down approach that starts from the existing genome to simplify or reduce the number of genes. While it may sound like a simple task, the challenge lies in deciding which genes are essential to keep and which to delete [29]. Since the end of the last century, the minimal genome has been implemented in bacteria through comparative approaches, which identify shared essential genes by comparing different species [28] and experimental approaches, which determine gene essentiality by systematically deleting genes [30]. Firstly, the comparative genomics approach relies on the hypothesis that genes conserved in distantly related species are likely essential for cellular function [28]. However, the limitation of this approach is that the same essential function can be performed by different non-orthologous genes. Secondly, the experimental gene inactivation approach relies on identifying genes that cause lethality when inactive. This could be achieved through massive transposon mutagenesis strategies [31, 32], the use of antisense RNA [33], or systematic activation of each gene in the genome [34]. The genome of eukaryotes is inherently more complex than that of prokaryotes, presenting significant challenges in minimizing it through conventional approaches. Trimming the genome of *S. cerevisiae* holds considerable appeal due to its diverse applications in research and industry. With 6,000 genes, 80% of which are deemed non-essential, totalling 12 Mbp in length, distributed across 16 chromosomes, *S. cerevisiae* offers an enticing target. Recent studies have demonstrated that deleting 85% of these non-essential genes can maintain over 90% of the wild-type phenotype [35, 36]. However, given that 90% of genes interact with each other, indiscriminate removal can lead to synthetic lethal interactions. Thus, a minimal genome must encompass essential genes and minimal gene sets to sustain life under specific conditions. Predicting which non-essential genes should accompany essential ones is challenging due to the genome’s complexity and the scattered distribution of essential genes across chromosomes [37, 38]. Despite these challenges, leveraging tools like SCRaMbLE holds promise for minimizing the synthetic genome efficiently for eukaryotic cells [39, 40]. Nonetheless, drawbacks for SCRaMbLE such as random genomic rearrangements and limitations in terms of deleting essential genes, particularly when associated with LoxPsym units, pose challenges [41]. We term LoxPsym units harbor essential genes ‘essential rafts,’ which can be relocated within the synthetic genome but cannot be removed due to their essentiality [42]. With the Sc2.0 project nearing completion, establishing an efficient workflow for minimizing the synthetic genome is crucial for advancing minimal genome research in higher eukaryotes. Previously, Wang et al. (2020) constructed strains harboring essential gene arrays in different configurations and observed minimal effects on gene expression. They applied SCRaMbLE to investigate synthetic lethal interactions in synIII (Wang et al., 2020). In addition, the Dai group compacted the arm of synthetic chromosome XIIL by deleting 39 of 65 nonessential genes using SCRaMbLE (Luo, Yu, et al., 2021). Together, these studies establish SCRaMbLE as a powerful approach for genome minimisation, enabling systematic identification of essential regions and large-scale genome reduction. However, defining minimal functional genomes while maintaining cellular fitness remains challenging. Here, we engineered neo-chromosome III variants harboring essential genes, utilising both native and refactored orthogonal regulatory elements sourced from diverse yeast species, including the closely related *S. paradoxus* and the distantly related *S. eubayanus*. This design increases sequence divergence, enhances stability, and enables systematic investigation of genome minimisation and cellular fitness. We then applied complementary strategies to trim synthetic chromosome III: CRISPR/Cas9 enables precise deletion of individual non-essential genes and is effective at small scales, whereas SCRaMbLE drives larger-scale genome reductions. However, synthetic lethal interactions constrain the extent of deletions, highlighting a key limitation in genome minimisation.

## Results

### 1. Characterization of promoter activity of synthetic essential genes harboring native and orthogonal regulatory elements derived from *Saccharomyces* species

Promoters play a central role in regulating gene expression in metabolic pathways and genetic circuits in *S. cerevisiae* [43]. Although approximately 876 endogenous promoters of *S. cerevisiae* have been characterized for genetic engineering applications (Keren et al., 2013), native promoters in *S. cerevisiae* present several limitations, including an insufficient number of well-characterized regulatory elements, limited dynamic range, and inadequate orthogonality to endogenous regulatory networks. The reuse of identical promoters for multiple genes in heterologous pathways has been identified as a major contributor to synthetic pathway instability (Peng et al., 2018). Previous studies have shown that *GAL* promoters characterised across different *Saccharomyces* species display a range of expression strengths and induction ratios comparable to those of the native *GAL* promoter in *S. cerevisiae* [44]. Building on this framework, we engineered alternative configurations of an artificial essential neo-chromosome in *Saccharomyces cerevisiae*. We first constructed a neo-chromosome carrying essential genes under native regulatory control. Subsequently, we evaluated strategies to enhance chromosomal stability, expand the incorporation of synthetic yet biologically compatible sequences, and assess the impact of orthogonal promoters on essential gene function and cellular fitness [45]. To this end, we refactored promoters of essential genes using orthologous regulatory elements from related *Saccharomyces* species. Given that promoter sequence divergence is a primary determinant of cis-regulatory variation, this approach enables systematic interrogation of how regulatory differences influence gene expression and fitness in a synthetic genomic context. We selected *S. paradoxus* as a closely related species (∼5 million years divergence) and *S. eubayanus* as a more distantly related member of the *Saccharomyces sensu stricto* clade, thereby establishing a gradient of regulatory divergence while preserving functional compatibility [46, 47]. Notably, *S. eubayanus* promoters combine reduced sequence homology with *S. cerevisiae* regulatory elements and the capacity to drive comparable expression dynamics, making them attractive components for genome engineering [48]. Furthermore, incorporation of divergent promoter sequences is expected to enhance neo-chromosome stability by limiting ectopic recombination. As homologous recombination is strongly dependent on sequence identity, reduced homology between orthologous promoters can suppress recombination through mismatch-repair–mediated anti-recombination mechanisms [49].To characterize promoter activity, all promoters were cloned as standardized biological parts into the bacterial HCKan-P vector using Golden Gate assembly with yeastFAB technology [50, 51], generating three promoter libraries. Each library contained 14 promoters of essential genes derived from *S. cerevisiae*, *S. paradoxus*, or *S. eubayanus*, respectively. Promoter activity was evaluated using a YFP/mCherry dual-reporter system to compare native and orthogonal promoters associated with essential genes on chromosome III. Promoters were subcloned upstream of YFP in a centromeric shuttle vector containing a dual-reporter cassette using Golden Gate assembly. The resulting 32 promoter constructs were individually transformed into wild-type *S. cerevisiae* BY4742. Cells were grown in triplicate in SC–Leu medium, diluted to an OD₆₀₀ of 0.1, and analysed using a plate reader (Fig. 1A). Promoter sequences of chromosome III essential genes were obtained from *S. cerevisiae* and compared with homologous sequences from the closely related species *S. paradoxus* and the more distantly related species *S. eubayanus* [52] (Fig. 1B). Promoter activity was quantified as the ratio of YFP to mCherry fluorescence. Relative activity of native promoters was normalised to the standardised promoter *CYC1p* and OD₆₀₀. Our results show that most *S. cerevisiae* promoters of essential genes on chromosome III function as weak promoters with minimal expression levels, with the notable exception of *PGK1p*, which exhibits expression comparable to the strong constitutive promoter *TDH3p* (Fig. 1C). We further analyzed cross-species differences in promoter strength between *S. paradoxus* and *S. cerevisiae*, and calculated the ratio of orthogonal promoter activity relative to their *S. cerevisiae* counterparts. A ratio of 1 indicates equivalent activity, values below 1 indicate reduced activity, and values above 1 indicate increased activity relative to *S. cerevisiae*. Each promoter was assayed at 8, 12, and 16 hours under aerobic conditions. Promoters *KKR1p1*, *PGS1p7*, and *RSC6p10* from *S. paradoxus* initially displayed more than twofold increase in relative activities compared to *S. cerevisiae*, but these differences diminished over time. In contrast, *CTR86p11* showed lower initial activity after 8h, but the activity increased over time, while the remaining *S. paradoxus* promoters exhibited activity similar to their *S. cerevisiae* counterparts. Similarly, we analyzed cross-species differences in promoter strength between *S. eubayanus* and *S. cerevisiae*, calculating the ratio of orthogonal promoter activity relative to their *S. cerevisiae* counterparts. In *S. eubayanus*, *PDI1p4* maintained a consistent twofold higher relative activity, whereas *PBN1p3* began with low activity after 8h, but it increased over time to reach 1. The rest of orthogonal promoters from *S. eubayanus* showed activity comparable to *S. cerevisiae* promoters (Fig. 1D–E). Sequence homology analysis showed that *S. cerevisiae* promoters are more similar to those of the closely related species *S. paradoxus* than to those of the more distantly related *S. eubayanus*. Promoter sequence divergence was 17.2% between *S. cerevisiae* and *S. paradoxus*, compared with 39.3% between *S. cerevisiae* and *S. eubayanus* (Fig. 1F). This pattern is consistent with established phylogenetic relationships within the *Saccharomyces* genus and indicates substantial conservation of promoter architecture between closely related species, with markedly increased divergence observed at greater evolutionary distances. Despite variation in orthogonal promoter activity, we proceeded to investigate how altered expression of essential genes driven by native or orthogonal promoters in essential new chromosome III affects essential gene functionality. These findings provide the foundation for engineering multiple versions of essential neo-chromosomes.

**Figure 1.**
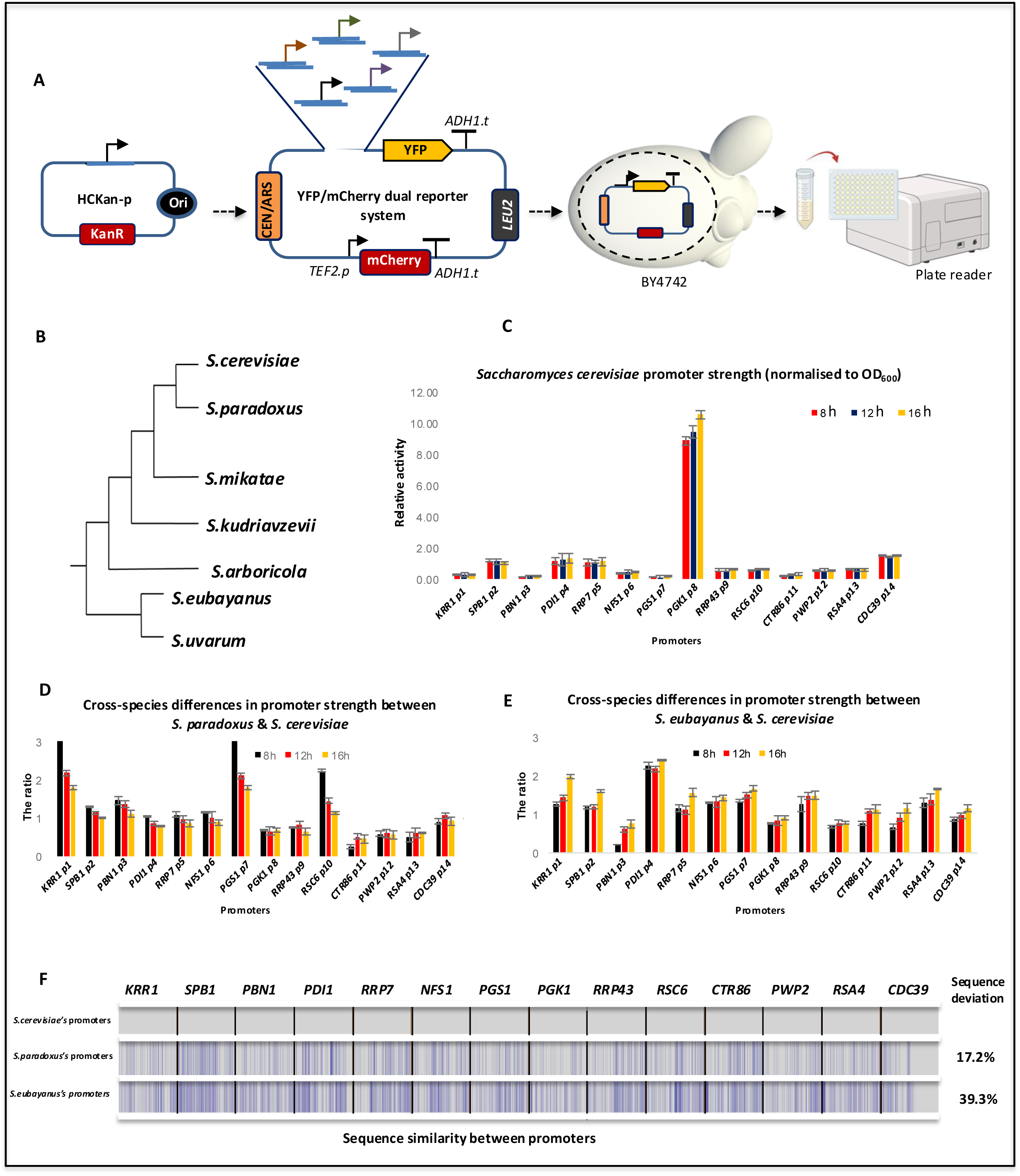
Orthogonal promoters derived from *S. paradoxus* and *S. eubayanus* exhibited levels of relative activity similar to those of the native promoters of essential genes on chromosome III of *S. cerevisiae*. **A.** Workflow used to characterize promoters: on the left side, all promoters were cloned into the bacterial HCKan-P vector using the Golden Gate (GG) reaction with yeastFAB technology and then subcloned into a shuttle centromeric vector containing the YFP/mCherry dual-reporter system, upstream of YFP, using a second GG reaction. The YFP/mCherry dual-reporter system was employed to measure promoter activity of native and orthogonal promoters of essential genes on chromosome III. The dual-reporter constructs harboring the tested promoters were individually transformed into wild-type *S. cerevisiae* BY4742. Three biological replicates of *S. cerevisiae* containing dual-reporter constructs harboring the tested promoters were grown overnight in SC-Leu medium, back-diluted to OD₆₀₀ = 0.1 and subjected to plate reader analysis. Promoter activity was defined as the ratio between YFP and mCherry fluorescence intensity, and relative activity was defined as the promoter activity normalized to *CYC1*p and OD₆₀₀. Fluorescence was measured using a Tecan plate reader with YFP detected at 514 nm excitation and 535 nm emission, and mCherry detected at 587 nm excitation and 610 nm emission. **B.** A phylogenetic tree of *S. cerevisiae* showing the closely related species *S. paradoxus* and the more distantly related species *S. eubayanus [52]*, from which promoter sequences were obtained. **C.** The schematic diagram illustrates the relative activity of native promoters of essential genes on chromosome III in *S. cerevisiae* normalized to *CYC1*p and OD₆₀₀. All *S. cerevisiae* promoters show weak promoter activity except for *PGK1*p, which exhibited a high level of expression. **D–E.** Graphs depict the ratio of relative promoter activity/OD₆₀₀ of orthogonal promoters from *S. paradoxus* and *S. eubayanus*, respectively, normalised to their *S. cerevisiae* counterparts. Promoter activity was determined after 8, 12, and 16 hours under aerobic conditions using a Biotek plate reader at 30 °C, and error bars represent ± SEM from three biological replicates. The relative activity of *KKR1*p1, *PGS1*p7, and *RSC6*p10 derived from *S. paradoxus* was initially >2 fold but decreased over time compared to their *S. cerevisiae* counterparts, whereas *CTR86*p11 showed lower initial activity that increased over time. The ratio of promoter activity of *PDI1*p4 derived from *S. eubayanus* was approximately 2, whereas *PBN1*p3 started at a low level after 8 h and increased over time. The remaining orthogonal promoters exhibited approximately promoter activity similar to their *S. cerevisiae* counterparts. Error bars are ± SEM from three biological replicates. **F.** The schematic diagram shows the sequence alignment between native promoters of essential genes on chromosome III in *S. cerevisiae* and orthologous promoters from *S. paradoxus* and *S. eubayanus*. Each promoter region is 500 bp; the first line includes 14 promoters from *S. cerevisiae*. Blue lines represent nucleotide mismatches between species, whereas white regions indicate conserved sequences. The sequence divergence in *S. eubayanus* promoters was substantially higher than in *S. paradoxus* promoters.

### 2. Evaluating the functionality of synthetic essential genes in the eNeo-chromosome constructed with native and orthogonal regulatory elements through yeast-species complementary assay

Previous studies have extensively investigated gene function using the comprehensive *Saccharomyces* gene deletion (YKO) collection. This resource contains isogenic knockouts for nearly all annotated yeast ORFs, with each ORF replaced by a kanMX4 cassette conferring resistance to G418 in BY4741 and BY4742 haploid strain [35]. Complementary approaches include conditional temperature-sensitive (TS) mutants, which carry point mutations that maintain gene function under permissive conditions but lose function under nonpermissive conditions [53]. A limitation of TS mutants is residual gene activity under nonpermissive conditions, which can obscure phenotypic interpretation. To overcome these limitations, we generated three libraries of shuffled strains to evaluate the functionality of essential genes expressed under native and orthogonal regulatory elements. A ‘shuffled strain’ is defined as a strain in which a resident plasmid is replaced by an alternative plasmid carrying a different selectable marker, enabling plasmid exchange through selection. Loss of the original plasmid was confirmed by the inability of the shuffled strain to grow on selective media corresponding to the marker carried by the initial plasmid. Each shuffled strain harbors a single essential gene (Eg) deletion in synthetic chromosome III and carries the corresponding Eg transcriptional unit (TU) assembled with native regulatory elements from *S. cerevisiae* or orthogonal regulatory elements from *S. paradoxus* or *S. eubayanus* on a low-copy pRS centromeric vector named pRS-eNeochromIII.V1-G418 (Fig. 2A). We used the SynIII strain as the genetic background because it serves as our primary platform for genome minimization studies. To construct shuffled strains, we started with the pRS–eNeochromosome III.v1-G418 SynIII strain and performed individual deletions of essential genes. The eNeo-chromosome III.v1 was then shuffled with POT2–*LEU2* plasmids carrying the corresponding essential genes, eliminating the need to repeat deletions for different regulatory contexts. This approach generated three libraries: (i) SynIII strains complemented with native *S. cerevisiae* essential genes assembled as eTUs with their native regulatory elements; (ii) shuffled strains complemented with orthologous genes assembled with regulatory elements from *S.paradoxus*; and (iii) equivalent strains complemented with orthologous genes assembled with regulatory elements from *S. eubayanus*.. Correct construction was verified by sequencing (described in a later section). Each essential gene on chromosome III was then individually deleted using homologous recombination with a *URA3* deletion cassette flanked by 80 bp homology arms targeting the gene of interest (Fig. 2C). Positive transformants lacking the essential gene from synthetic chromosome III were selected on SC-Ura medium and verified by confirmation PCR targeting upstream and downstream chromosomal integration sites. Subsequently, POT2-*LEU2* vectors carrying the corresponding essential gene (POT2-*LEU2*-Eg), which has been deleted in chromosome III, assembled with either native or orthogonal regulatory elements, were individually co-transformed into SynIII strains lacking the respective essential gene (Fig. 2B). Transformants harboring POT2-*LEU2*-Eg constructs were selected on SC-Leu medium and confirmed by PCR. To obtain the final shuffled strains, cultures were incubated in SC-Leu medium for 24 hours to maintain selection for POT2-*LEU2*-Eg while removing the G418 selection, thereby allowing for the loss of the pRS-eNeochromIII.V1-G418 construct. Successful shuffled strains were identified by their ability to grow on SC-Leu medium and their inability to grow on YPD + G418, confirming loss of the original eNeo-chromosome construct pRS-eNeochromIII.V1-G418. The resulting strains contain a single essential gene deletion in synthetic chromosome III complemented by an individual Eg transcriptional unit (eTU) on a centromeric vector. After constructing three libraries of shuffled strains, we evaluated their ability to complement essential gene deletions in synthetic chromosome III. Phenotypic analysis of the three shuffled strain libraries on selective SC-Leu medium demonstrated that plasmid-encoded essential genes compensated for the chromosomal deletions (Fig. 3A). However, complementation was associated with partial fitness defects in some strains, reflected by reduced colony size. For example, the *ctr86Δ* eNeochromIII.V1Δ SynIII+POT2-*CTR86* strain complemented with the *S. paradoxus* regulatory element displayed smaller colonies, consistent with the lower promoter activity observed for the orthogonal *CTR86* promoter. More broadly, most shuffled strains exhibited smaller colony size compared with the SynIII+pRS415 control strain, suggesting a modest fitness cost. This effect may arise from variation in essential gene dosage, as Eg are typically maintained at one copy per haploid genome, but it fluctuates to 2–5 copies on pRS vector. Deletion of *PDI1* from the SynIII background proved technically challenging using the *URA3* deletion strategy, preventing evaluation of its functionality using the standard shuffled-strain workflow. Incorporation of the native *PDI1* gene into the pRS-eNeochromIII.V1 did not support cell viability. This suggests that the DNA sequence targeted for removal may contain an unrecognized functional element. To address this limitation, we adopted an alternative strategy to assess *PDI1* constructs assembled with native or orthogonal regulatory elements. An engineered *Saccharomyces cerevisiae* BY4742 with WT background strain carrying an inactive chromosomal *PDI1* allele and harboring an orthogonal translation system (OTS) on a CEN/ARS centromeric vector to regulate *PDI1* expression served as the starting platform (Fig. 3B). In this strain, a synthetic noncanonical amino acid–dependent system controls *PDI1* expression through the introduction of an amber stop codon into the *PDI1* open reading frame via CRISPR/Cas9. In the absence of O-methyltyrosine (OMT), translation terminates prematurely, producing a nonfunctional truncated protein and rendering cells inviable without OMT supplementation. The OTS was maintained on a plasmid with *URA3* auxotrophic selection; Removal of URA3 selection pressure resulted in loss of the OTS plasmid, which encodes a suppressor tRNA that decodes the UAG codon and a dedicated synthetase that incorporates OMT, thereby enabling full-length *PDI1* translation. To assess functional complementation, we engineered *PDI1* transcriptional units containing regulatory elements from *S. cerevisiae*, *S. paradoxus*, or *S. eubayanus*. These constructs were assembled using Golden Gate and yeastFAB methods, cloned into the POT2-LEU2 vector, and individually transformed into the BY4742 OTS strain. Upon removal of *URA3 s*election while maintaining LEU2 selection, the OTS plasmid was lost, and cells retaining the POT2-LEU2 constructs were selected, as confirmed by replica plating (Fig. 2a). Cell viability was maintained by the POT2-LEU2–encoded *PDI1* constructs in the BY4742 background; however, colony sizes were modestly reduced, likely reflecting variation in essential gene dosage. Notably, *PDI1* constructs assembled with both native and orthogonal regulatory elements successfully complemented loss of the chromosomal gene. These findings are consistent with observations from shuffled strains derived from the SynIII synthetic background, supporting the conclusion that orthogonal regulatory elements can sustain essential gene function, albeit with minor fitness trade-offs.

**Figure 2.**
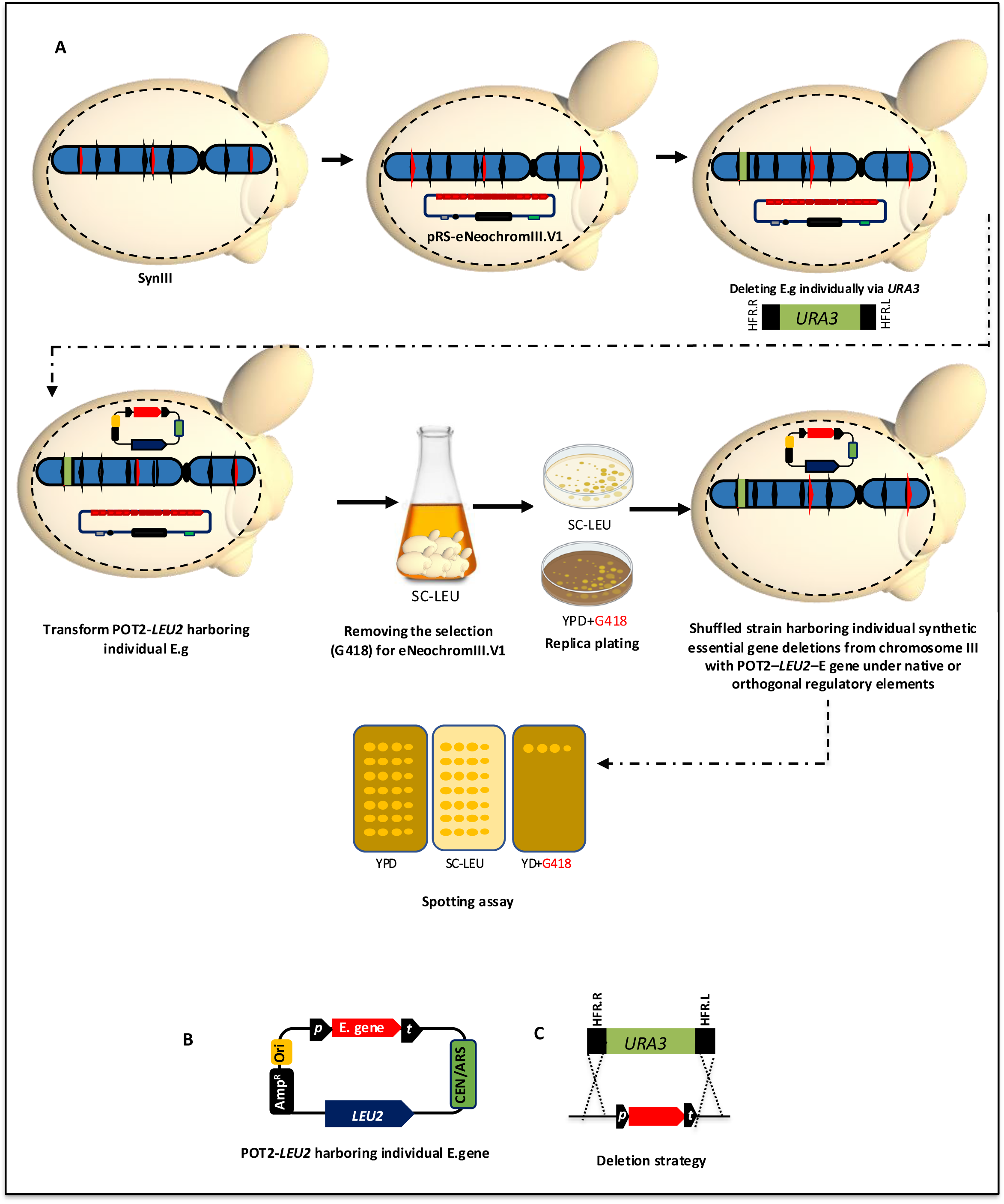
Workflow for engineering a shuffled SynIII strain library to assess the functionality of constructed essential genes regulated by native or orthogonal elements. **A.** Strategy to engineer the shuffled strain library: each shuffled strain harbors a single essential gene (Eg) deletion in synthetic chromosome III and carries an individual Eg transcriptional unit (TU) constructed with native (*S. cerevisiae*) or orthogonal regulatory elements from *S. paradoxus* and *S. eubayanus* on a pRS centromeric vector. First, pRS-eNeochromIII.V1 containing 14 essential genes constructed with native regulatory elements was transformed into the SynIII strain to generate the pRS-eNeochromIII.V1-G418 SynIII strain, which was confirmed by sequencing. This strain served as the starting platform. Each essential gene on synthetic chromosome III was individually deleted using homologous recombination with a URA3 deletion cassette. Synthetic mutants lacking the respective Eg were selected on SC-URA minimal medium and verified by PCR. Subsequently, the POT2-LEU2 vector carrying the corresponding Eg, matching the gene deleted from synthetic ChrIII, was co-transformed into the strain lacking that Eg. Transformants harboring the POT vector with the Eg constructed with native or orthogonal regulatory elements were selected on SC-Leu medium. To remove pRS-eNeochromIII.V1-G418 carrying the 14 essential genes with native regulatory elements, selective pressure was withdrawn by cultivating the strains on SC-Leu without G418. Transformants were plated on SC-Leu and replica-plated onto YPD+G418. The appropriate shuffled strains, containing a single Eg deletion in synthetic chromosome III and one individual Eg TU on a pRS CEN/ARS centromeric vector, were confirmed by growth on SC-Leu and inability to grow on YPD+G418. A spot assay was then performed on all shuffled strains to evaluate fitness and to confirm removal of pRS-eNeochromIII.V1-G418. **B**. The POT CEN/ARS centromeric vector vectors harbored essential genes constructed with native or orthogonal regulatory elements. **C**. Essential gene deletion strategy using the URA3 cassette which contains (p, ORF, T) with homologous flanking regions (HFR, 80 bp).

**Figure 3.**
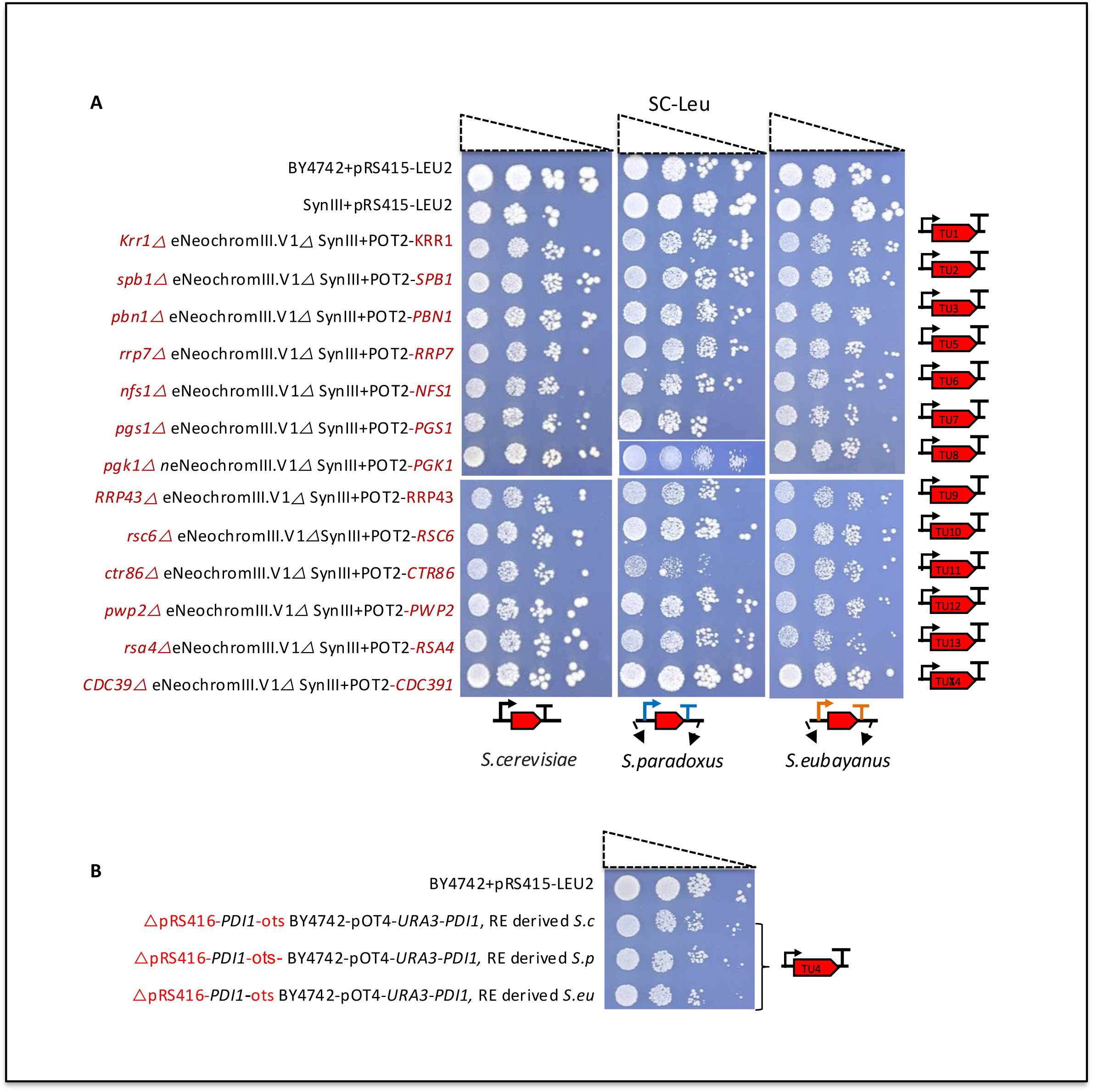
Essential genes engineered with native or refactored regulatory elements in different eNeoChromosome versions compensate for the absence of the corresponding essential genes on chromosome III. **A.** Phenotypic analysis of shuffled strain libraries to assess the fitness of essential genes engineered with native or refactored regulatory elements. Phenotypic analysis was performed on three libraries of shuffled strains grown on selective medium (SC-Leu). Each library contained 13 shuffled strains to characterize the functionality of essential genes engineered with native or refactored regulatory elements, except *PDI1*. In each shuffled strain, a single essential gene (Eg) was deleted from synthetic chromosome III (synChrIII) and complemented by the corresponding Eg engineered as a transcriptional unit (TU). These TUs were constructed either with native regulatory elements (REs) from *Saccharomyces cerevisiae* (left column), orthogonal REs from the closely related species *S. paradoxus* (middle column), or from the more distantly related species *S. eubayanus* (right column). Native or refactored essential genes were expressed from a (POT2-*LEU2*) CEN/ARS centromeric vector. All shuffled strains harboring Eg constructs (with either native or orthogonal REs) compensated for the absence of the corresponding chromosomal Eg through plasmid-encoded expression from the POT2-*LEU2* CEN/ARS centromeric vector. Most shuffled strains exhibited smaller colony sizes compared with the control strain SynIII+pRS415-*LEU2*, suggesting a mild fitness burden. This burden is likely caused by increased gene dosage, as essential genes normally present as a single copy in the haploid strain while in the shuffled strains, the essential genes were maintained at approximately 2–5 copies on the centromeric vector (POT2-*LEU2*). For example, *PGK1* and *CTR86* in the second library, when regulated by *S. paradoxus*-derived REs, showed reduced colony sizes, whereas *CDC39* displayed colony sizes comparable to the control strain SynIII+pRS415-*LEU2* across all three libraries. **B.** Alternative approach for evaluating *PDI1* functionality engineered with native or orthogonal regulatory elements. *PDI1* could not be deleted from synthetic ChrIII in SynIII using the standard deletion strategy employed to generate the shuffled strain libraries. The parental strain BY4742 + POT4-*URA3*–OTS-*PDI1* was used as the starting point to generate shuffled strains. This strain, has a wild-type background, already contained an orthogonal translation system (OTS) regulating *PDI1* expression within the POT4-*URA3*–OTS-*PDI1* cassette and harbored an inactive *PDI1* allele at the native chromosomal locus. *PDI1* was engineered using RE derived from *S. cerevisiae* (Sc), *S. paradoxus* (Sp), and *S. eubayanus* (Se), and each construct was cloned into the POT2-*LEU2* destination vector using Golden Gate and YeastFab technologies. These three constructs were individually transformed into the BY4742 strain carrying the OTS for *PDI1*. Following removal of URA selection pressure, loss of the OTS for *PDI1* was confirmed by replica plating, as strains were unable to grow on SC-Ura medium but survived on SC-Leu medium due to the presence of *PDI1* encoded on POT2-*LEU2*. Although colony sizes were slightly reduced, likely due to variation in essential gene dosage in the resulting strains, *PDI1* constructs containing native and orthogonal REs effectively complemented the absence of the native chromosomal *PDI1*. These results are consistent with previous observations obtained using shuffled strains derived from the SynIII synthetic strain background.

### 3. Design and engineering strategy for the Essential Neo-chromosome version I (pRS-eNeochrome III.V1 *S.c*)

Synthetic yeast chromosomes and tRNA neochromosomes are commonly constructed using the SwAP-In (Switching Auxotrophic Progressively by Integration) strategy, in which DNA mini-chunks are sequentially assembled via homologous recombination with iterative marker exchange (Annaluru et al., 2014; Schindler et al., 2023). In contrast, the Essential Neo-chromosome III.V1 (eNeochrome III.V1) was engineered using a non-iterative assembly framework designed to preserve native regulatory architecture while enabling systematic reorganization of essential gene content. Version I of eNeochrome III (PRS-eNeochrome III.V1 *S.c*) was assembled *in vivo* in a SynIII synthetic yeast background using endogenous homologous recombination machinery. This design enables controlled investigation of essential gene organization, genomic architecture, and synthetic genetic interactions. All essential genes were arranged in a unidirectional tandem configuration independent of their native chromosomal orientation, allowing assessment of positional and orientation constraints on essential gene function. eNeochrome III.V1 *S.c* contains the 14 essential chromosome III genes in the following order: *KRR1, SPB1, PBN1, PDI1, RRP7, NFS1, PGS1, PGK1, RRP43, RSC6, CRTR86, PWP2, RSA4,* and *CDC39*. The essential tRNA gene *SUP61* was excluded because its function is supplied by the tRNA neochromosome (Schindler et al., 2023) in the final Sc2.0 strain comprising 16 synthetic chromosomes plus the tRNA neochromosome. In the SynIII strain, *SUP61* is integrated at the HO locus as part of chromosome III minimization. Each essential gene was constructed as an independent transcriptional unit comprising a 500 bp native promoter, the open reading frame, and a 200 bp terminator. To enable future programmable genome rearrangements, a DRE/rox site-specific recombination system was incorporated, with each transcriptional unit flanked by 32 bp rox sites positioned downstream of the terminator (Fig. 4A). Mini-arrays containing four or five transcriptional units were assembled in centromeric pRS vectors carrying distinct selectable markers. Wild-type genomic DNA from strain BY4742 served as the PCR template to preserve PCRTag distinguishability between wild-type-derived essential genes and their synthetic chromosome counterparts [54]. WT PCRTags verified essential gene incorporation into the eNeochromosome III, whereas Syn PCRTags confirmed retention of essential loci in the synthetic chromosome III background. Assembly proceeded in three stages. First, each essential gene was PCR-amplified with 60 bp homology flanking regions to enable *in vivo* homologous recombination. Three mini-essential arrays were independently assembled in the SynIII strain: (i) *KRR1, SPB1, PBN1, PDI1 and RRP7* in pRS413 (*HIS3*), (ii) *NFS1, PGS1, PGK1, RRP43* and *RSC6* in pRS415 (*LEU2*), and (iii) *CRTR86, PWP2, RSA4* and *CDC39* in pRS416 (*URA3*). Each mini-array was flanked by unique restriction sites (*Sbf*I, *Mlu*I, and *Srf*I, respectively), enabling *in vitro* excision of the mini-arrays and facilitating their subsequent *in vivo* assembly into the eNeochromosome III. A unique sequence of 100bp flanked both end of the eNeochromosome to allow excision and hierarchical assembly, enabling future scaling toward larger essential megachromosomes harboring 1000 essential genes from across the genome for SC3.0 project. Correct assembly of each mini-array was confirmed by junction PCR and WT PCRTag analysis. Validated mini-arrays were recovered from yeast and propagated in bacteria using the EASY-C protocol (REF), followed by restriction digestion to release purified mini-arrays. In the final stage, the three mini-arrays were assembled *in vivo* into the complete eNeochromeIII.V1 *S.c* via homologous recombination machinery in SynIII synthetic strain (Fig. 4C). Final chromosome assembly was verified by junction PCR and WT PCRTag analysis (Fig. 4D). The assembled eNeochromosome in SynIII strain was subsequently recovered using the EASY-C strategy [55]. The observed restriction digest profiles matched the predicted restriction pattern (Fig. 4E). Linearised high-quality DNA was obtained via the EASY-C strategy, and the sequence was confirmed by long-read nanopore sequencing, validating structural integrity of the engineered eNeochromosome (Fig. 4F).

**Figure 4.**
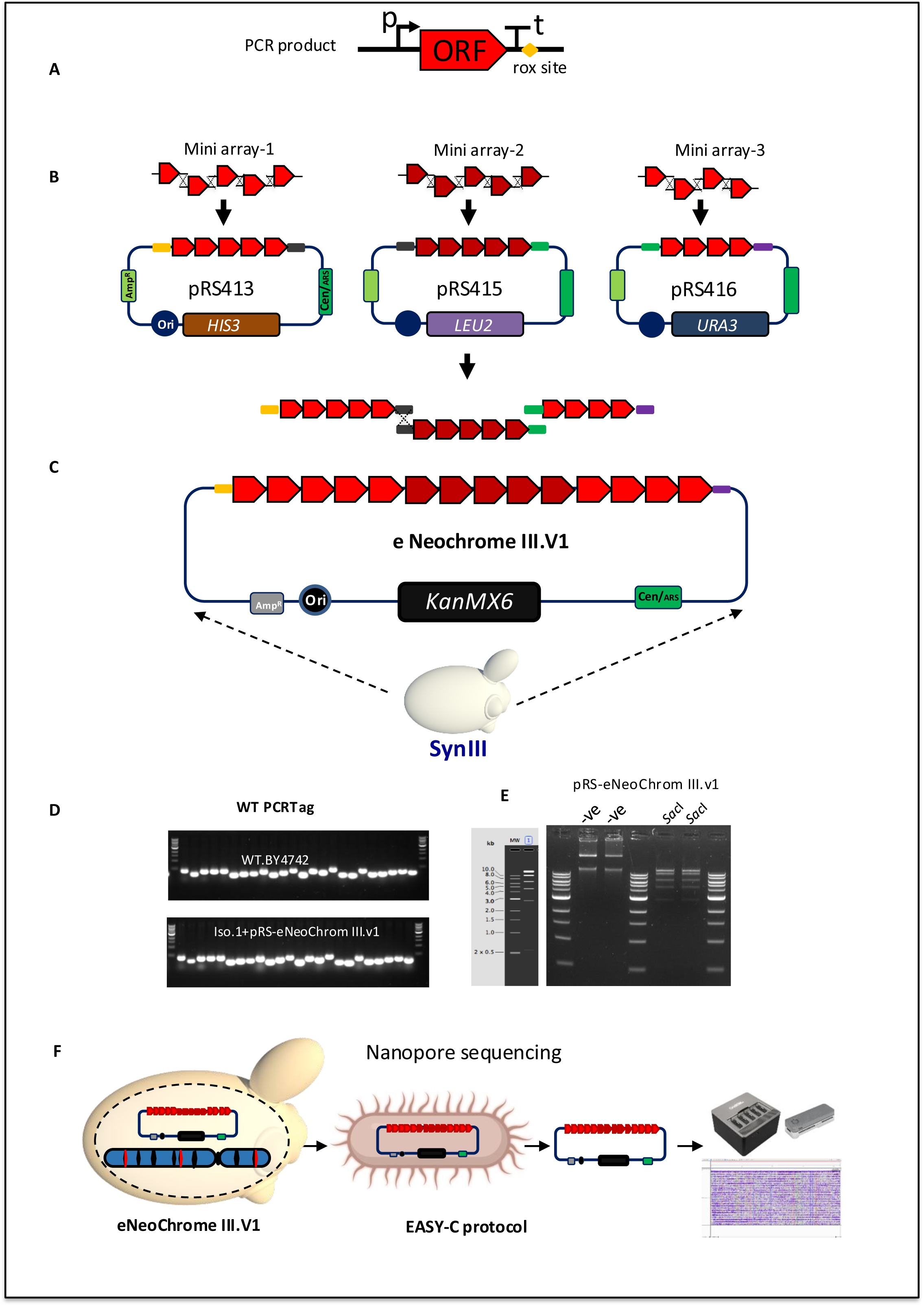
Design and construction strategy for engineering eNeochrome III.V1. **A.** Schematic diagram of the transcriptional unit (TU) for essential genes, consisting of a 500 bp promoter (p) upstream of the open reading frame (ORF), ORF, followed by a 200 bp terminator (t) downstream of the ORF, with a ROX site positioned downstream of the terminator. Although the serine tRNA gene *SUP61* (tRNA-Ser) is an essential gene, it is not included in this design, as it will be incorporated into the tRNA neochromosome, which harbors all tRNA genes in the final synthetic strain. Each TU was PCR-amplified from genomic DNA of the wild-type (WT) strain BY4742, with 60 bp homology-flanking regions (HFRs) to adjacent TUs or the plasmid backbone. **B.** Schematic illustration of the construction strategy for assembling three essential mini-arrays via homologous recombination machinery (HR) *in vivo* in yeast. The first mini-array contains five essential genes (Egs) cloned into pRS413-*HIS3*, the second contains five Egs in pRS415-*LEU2*, and the third contains four Egs in pRS416-*URA3*. All Egs are arranged unidirectionally and in a defined order, regardless of their native genomic orientation. Each mini-array is flanked by unique restriction enzyme sites designed to release the array *in vitro*. Each mini-array was assembled individually *in vivo* using the SynIII synthetic strain. Successful assembly was confirmed by junction PCR in combination with WT PCRTags. Following confirmation, the pRS plasmids harboring the essential mini-arrays were extracted using the EASY-c protocol and digested with restriction enzymes to release the essential mini-arrays. **C.** Schematic diagram illustrating the assembly of the three essential mini-arrays into a single construct via homologous recombination using pRS-KanMX6 as the destination vector *in vivo* in the SynIII strain. **D.** Successful assembly of eNeochrome III.V1 was confirmed using WT PCRTags, with the WT strain BY4742 serving as a positive control. **E.** eNeochrome III.V1 was extracted by EASY-c protocol and analysed by restriction digestion, revealing the expected digestion pattern. **F.** The EASY-c protocol was used to extract eNeochrome III.V1 from the SynIII strain, propagate it in bacteria, purify and linearize the eNeochrome III.V1 construct, and subject it to Nanopore sequencing. Nanopore sequencing analysis confirmed the correct assembly of eNeochrome III.V1.

### 4. Introducing an orthogonal recombination system to modify copy number variation in eNeochrome III.V1 *S.c*

Our design extends this framework by incorporating rox-based programmable recombination sites combined with modular hierarchical assembly, establishing a scalable platform for independent SCRaMbLE analysis of the eNeochrome V1 *S.c*. A site-specific recombination system based on Dre/rox was introduced into eNeochrome III.V1 *S.c.* Each transcriptional unit is flanked by 32 bp rox recombination sites positioned downstream of the transcriptional terminator **(Fig. 4A).** The Dre/rox system is orthogonal to Cre-lox. Previous studies have demonstrated the absence of cross-reactivity between Dre/rox and Cre-lox recombination systems; consequently, Dre-mediated recombination does not interfere with SCRaMbLE events targeting synthetic chromosome III [56]. This architecture provides an additional degree of regulatory control, enabling eNeochrome III.V1 *S. cerevisiae* to undergo SCRaMbLE independently of the synthetic chromosome and allowing controlled modulation of eNeochrome III.V1 copy number to support downstream functional analyses. This system presents new opportunities for future experimental exploration and functional characterisation.

### 5. Refactoring regulatory elements of essential chromosome III genes using YeastFab

Refactoring the regulatory elements (RE) of essential chromosome III genes using conventional cloning approaches would be labor-intensive and difficult to standardize. To overcome this limitation, we employed YeastFab technology, a modular assembly platform that enables rapid construction of metabolic pathways from standardized biological parts, including promoters, open reading frames (ORFs), and terminators [50, 51]. This framework was used to construct eNeochrome variants. V2 was built with native regulatory elements from *S. cerevisiae* while, V3 and V4 were built by replacing native regulatory elements from *S. cerevisiae* with orthogonal elements derived from the closely related species *S. paradoxus* and the more distantly related *S. eubayanus*, using YAC12 as the destination vector to generate both circular and linear forms. Andrew W. Murray and Jack W. Szostak described the construction of artificial chromosomes in yeast [57]. A key feature of yeast artificial chromosomes (YACs) is the presence of an autonomously replicating sequence (ARS), which enables extrachromosomal replication. In addition, YACs contain a centromere (CEN), such as CEN4, which ensures stable, single-copy propagation by improving segregation fidelity and reducing copy number variation. Telomeres, which are specialised DNA sequences at the ends of chromosomes, are also essential components required for the stability and maintenance of linear DNA. YACs are derived from bacterial vector systems and are used to clone large DNA fragments. They typically include three auxotrophic markers for selection. Furthermore, YACs can exist in both circular and linear forms; removal of the marker located between telomeric sequences allows conversion into the linear form. This strategy enables rapid and efficient swapping of regulatory elements and supports the construction of multiple synthetic eNeochrome variants, facilitating downstream stability assays and investigations into the effects of altered essential gene expression on cellular fitness and genome minimization. Figure 5 illustrates the overall refactoring workflow. Each component was PCR-amplified or synthesized and formatted as a standardized biological part (Fig. 5A). ORFs were cloned into HCKan-O vectors to generate an ORF library, promoters into HCKan-P vectors to generate promoter libraries, and terminators into HCKan-T vectors to generate terminator libraries (Fig. 5B). These parts are mutually compatible and enable one-pot Golden Gate assembly of complete TUs (Fig. 5C). Native regulatory elements from *S. cerevisiae* and orthogonal elements from *S. paradoxus* and *S. eubayanus* were assembled to investigate their effects on gene regulation, phenotype, cellular fitness, and genome minimization. We generated an ORF library comprising 14 essential chromosome III genes and three promoter libraries: native *S. cerevisiae* promoters and orthogonal promoters from *S. paradoxus* and *S. eubayanus*. Corresponding terminator libraries were constructed in parallel. To improve Golden Gate cloning efficiency, internal *Bsa*I and *Bsm*bI sites were removed from ORFs via codon optimisation. *Bsa*I was used for library construction in HCKan vectors, and *Bsm*bI for TU assembly into the POT-*URA3* destination vector. All parts were verified by restriction digest and Illumina sequencing. Fourteen TUs were temporarily subcloned into five pYFASS bacterial vectors (Fig. 5D), with each vector carrying three TUs except the final vector, which carried two. Correct constructs were verified and released by restriction digestion. The five purified fragments were assembled *in vivo* with YAC12 via homologous recombination in the synIII strain, generating circular or linear eNeochromes (Fig. 5E & F). Circular forms were produced by co-transforming linearized YAC12 with the five fragments, whereas linear forms were generated after removal of the *LEU2* marker between telomeres before transformation. Successful assembly was confirmed using WT PCRTags (Fig. 5G) and long-read nanopore sequencing directly from yeast genomic DNA. Interestingly, read depth was substantially higher for linear chromosomes (>2000 reads) compared to circular forms (66 reads) (Fig. 5H & I). Despite lower read counts relative to bacteria-purified circular chromosomes, nanopore sequencing provided sufficient coverage to validate assembly. Our previous work has shown that shuttle chromosomes can be extracted from yeast and propagated in bacteria via EASY-C protocol [55] to improve DNA yield and read depth as it avoids competition with homologous genomic sequences from yeast. Four eNeochrome variants incorporating native and orthogonal regulatory elements have been constructed (Fig. 6). Strains carrying these variants were prepared for comprehensive phenotypic, fitness, and chromosomal stability analyses across 100 generations.

**Figure 5.**
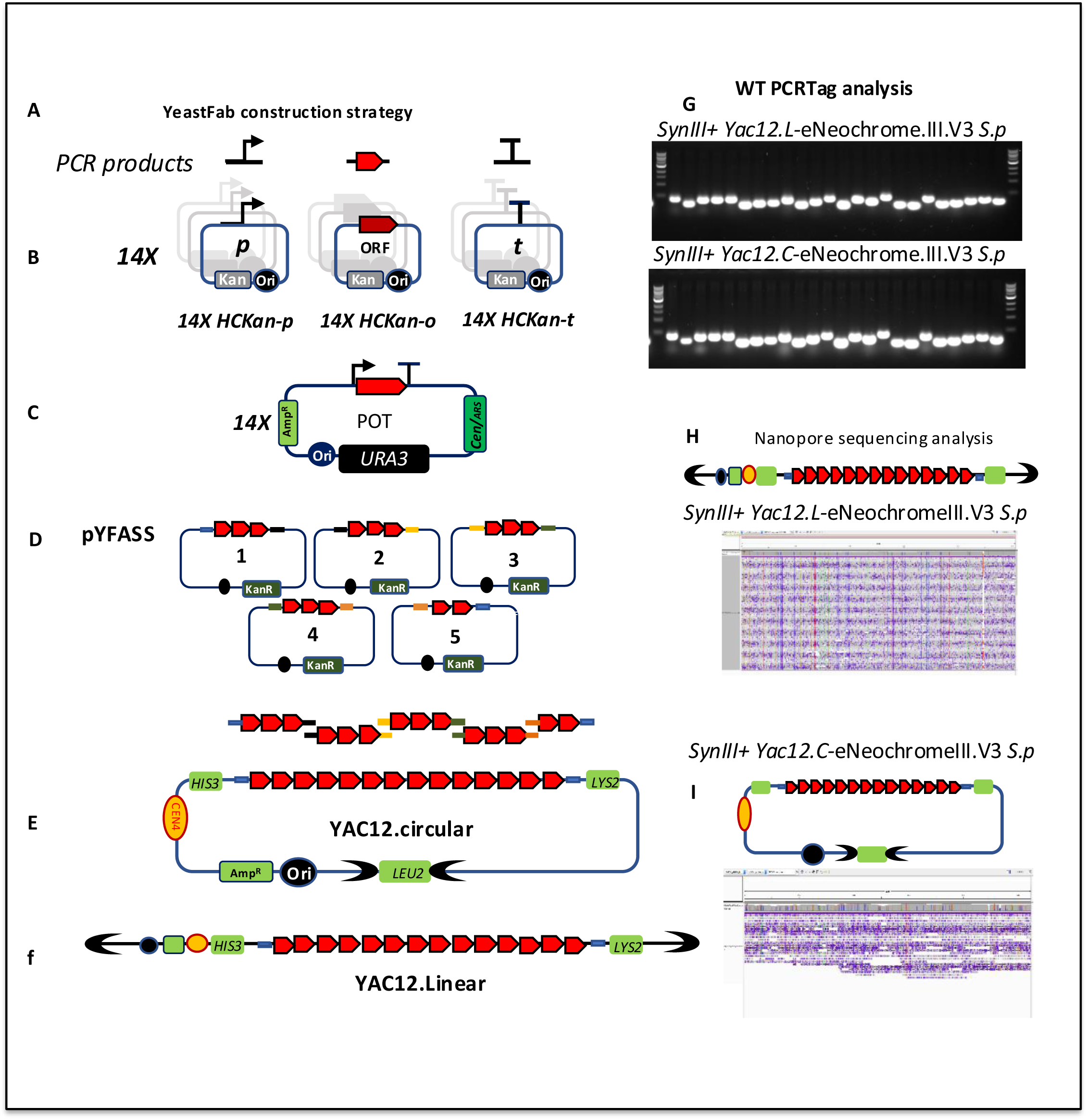
Design and construction strategy for refactoring regulatory elements of essential genes from *S. cerevisiae*, *S. paradoxus*, and *S. eubayanus* using YeastFab technology to generate eNeochrome III versions V2, V3, and V4 in linear and circular forms. Schematic overview of the strategy used to engineer different versions of eNeochrome III harboring essential genes (Egs) with native or orthogonal regulatory elements (REs), assembled in both linear and circular forms. **A.** Each standardized biological part of the transcriptional unit (TU)—promoter (p), open reading frame (ORF), and terminator (t)—was PCR-amplified or synthesized with compatible overhangs and cloned into the HCKan vector using Golden Gate (GG) assembly to generate promoter library, ORF library and terminator library. **B.** Promoters of essential genes located on chromosome III from three yeast species (*S. cerevisiae*, *S. paradoxus*, and *S. eubayanus*) were individually cloned into HCKan-p vectors to generate three promoter libraries, each containing 14 promoters. The first library contains native promoters from *S. cerevisiae*, while the second and third libraries contain orthogonal promoters derived from *S. paradoxus* and *S. eubayanus*, respectively. ORFs corresponding to the essential genes on chromosome III were derived from wild-type *S. cerevisiae* BY4741 and cloned individually into HCKan-O vectors to generate a library of 14 ORFs. Terminators of essential genes on chromosome III were cloned into HCKan-t vectors to form three terminator libraries (14 terminators each): native *S. cerevisiae* terminators and orthogonal terminators from *S. paradoxus* and *S. eubayanus*. **C.** Native or orthogonal regulatory elements were designed to be compatible with ORF parts from different libraries. Individual TUs were assembled in a single one-pot (POT) Golden Gate reaction using POT-*URA3* vectors. Newly constructed assemblies were verified by restriction digest analysis to release the insert using *Bsa*I. **D.** Two -three assembled essential Tus were sub-cloned into five YFASS vectors. The essential genes arranged in the same native order as in wild-type *S. cerevisiae* and correct assembly was verified by restriction digest analysis using *Bsm*BI. **E** Inserts were released and purified from the five YFASS vectors and flanked with 100-bp homologous flanking regions (HFRs) to facilitate in *vivo* assembly via homologous recombination (HR) in the SynIII strain to generate either linear or circular versions of eNeochrome III, using YAC12 as the destination vector. **F.** For linear constructs, the *LEU2* marker located between telomeres was removed prior to final assembly. **G.** Correct assembly of eNeochrome III was confirmed by wild-type PCRTag analysis. Both linear (YAC12.L-eNeochrome.V3) and circular (YAC12.C-eNeochrome) forms were validated (for instance PCRTag analysis here is for third version of the eNeoIII.V3 were RE derive from *S.paradoxus*). No product was observed for SynIII lacking eNeochrome III. **H.** Nanopore sequencing performed directly from yeast cells confirmed correct assembly of essential neochromosomes containing native or orthogonal regulatory elements for eNeochrome III variants (V2, V3, and V4). **I**. Linear constructs yielded substantially higher read coverage compared to circular constructs assembled with the same regulatory configurations.

**Figure 6.**
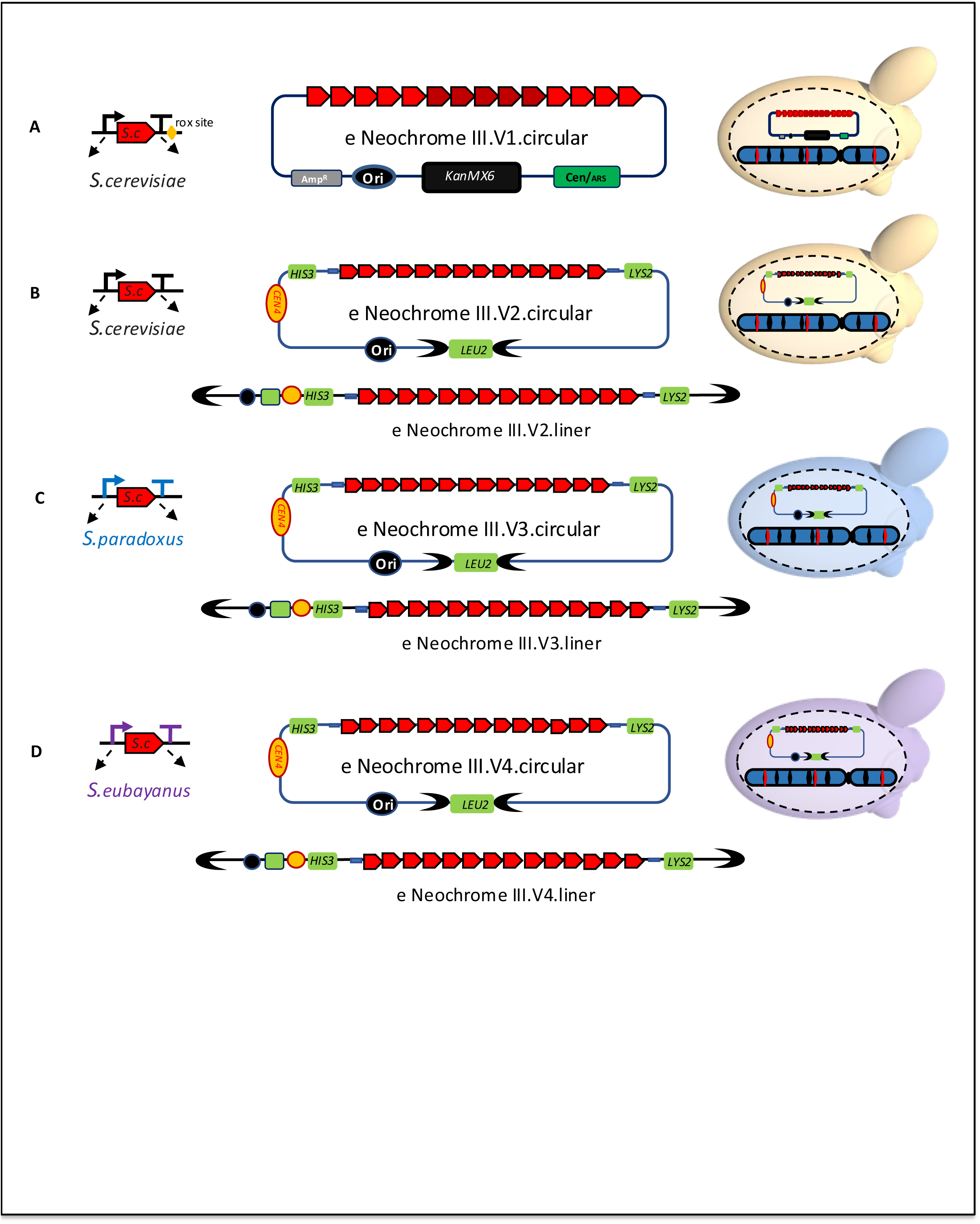
Different versions of eNeochrome III variants (V1–V4) constructed with native and orthogonal regulatory elements (REs). **A.** eNeochrome III variant V1, in which the REs of Egs are derived from *S. cerevisiae* and flanked by rox sites downstream of terminator. PRS vector, (ARS/ori, low-copy, ∼2-4 copies per cell) was used as the destination vector. **B.** eNeochrome III variant V2, in which the REs of Egs are derived from *S. cerevisiae* and lack rox sites downstream of the terminator. YAC12 was used as the destination vector to construct both linear and circular forms, typically present as a single copy. **C.** eNeochrome III variant V3, in which the REs of Eg are derived from *S. paradoxus*. YAC12 was used as the destination vector to generate both linear and circular forms, typically present as a single copy. **D.** eNeochrome III variant V4, in which the regulatory elements of essential genes are derived from *S. eubayanus*. YAC12 was used as the destination vector to generate both linear and circular forms, typically present as a single copy.

### 6. Comparative phenotypic analysis reveals robust growth in SynIII strains hosting YAC12-based eNeochromes, while pRS-based eNeochrome III.V1. *S.c.* causes severe growth defects

The SynIII+pRS-eNeochrome III.V1. *S.c.* strain exhibits impaired growth relative to both the SynIII and wild-type BY4742 strains in rich medium (YPD+G418) and minimal medium (SCD+G418) (Fig. 7A). Removal of the pRS-eNeochrome III.V1. *S.c.* construct restores normal growth, confirming that the plasmid is the primary driver of the phenotype. This defect is likely caused by essential gene dosage imbalance: essential genes are normally present as a single chromosomal copy, whereas expression from a low-copy centromeric pRS vector may result in 2–5 copies per cell, creating a burden on cellular homeostasis. An additional contributing factor may be the presence of 14 rox sites positioned downstream of essential genes, which could impose further cellular stress, as previously observed in tRNA neochromosome systems [19]. Growth curve analysis revealed a pronounced reduction in growth rate during logarithmic phase for the SynIII+pRS-eNeochrome III.V1. *S.c* strain, persisting for approximately 20 hours in liquid culture before stabilizing near stationary-phase levels comparable to SynIII and BY4742 controls (Fig. 7B). Notably, SynIII strains hosting all YAC12-based eNeochrome variants—containing either native regulatory elements from *S. cerevisiae* or orthogonal elements from *S. paradoxus* and *S. eubayanus*—consistently display robust phenotypes compared to the control strain SynIII+ empty YAC12. The YAC12 system maintains a single additional chromosomal copy of essential genes, which is more readily tolerated than plasmid-based overexpression. This is consistent with previous observations that yeast artificial chromosomes (YACs) segregate as single copies and exhibit mitotic stability in *Saccharomyces cerevisiae* haploid cells [58, 59]. This robustness is supported by both solid-media spot assays and 24-hour liquid growth analyses (Fig. 7C-F).

**Figure 7.**
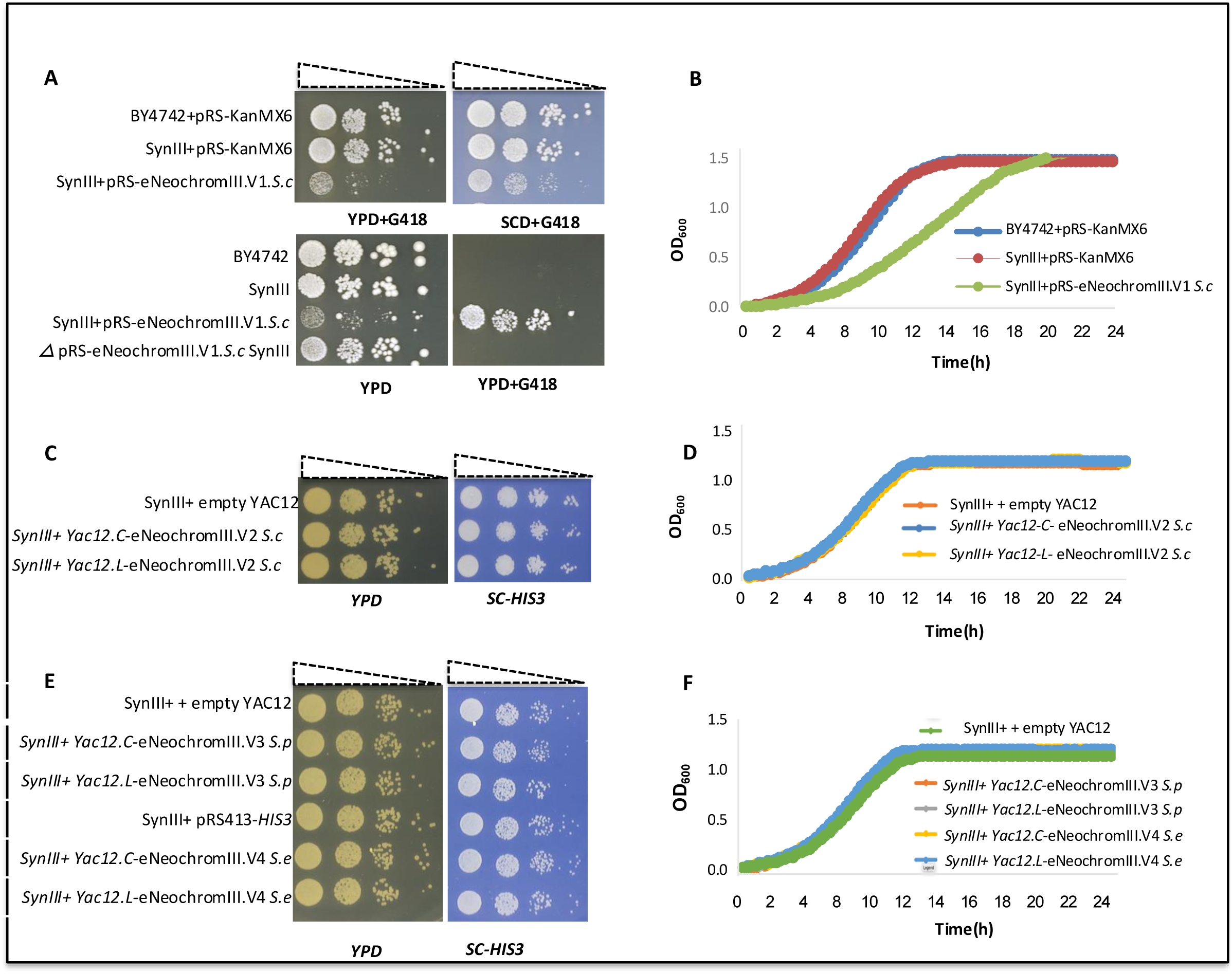
SynIII strains exhibit uniform growth phenotypes across eNeochrome III variants on YAC12, with the exception of eNeochrome III.v1, which causes a growth defect. **A.** SynIII cells expressing eNeochrome III.v1 from pRS show a pronounced growth defect relative to SynIII and WT BY4742 controls carrying the empty pRS-G418 vector on YPD+G418 and SCD+G418 media. Loss of the pRS–eNeochrome III.v1 plasmid restores normal growth in the Δ pRS–eNeochrome III.v1 SynIII strain. **B.** Growth curve analysis reveals a significant reduction in logarithmic-phase growth of the SynIII + pRS–eNeochrome III.v1 strain; however, after 20 h in liquid culture, cell density reaches levels comparable to those of SynIII and WT BY4742 in stationary phase. Error bars indicate ± SEM (n = 3 biological replicates). **C.** Spot assays and growth analyses show a consistent healthy phenotype for SynIII strains harboring eNeochrome III.v2 on YAC12 containing native recombination elements, in both linear (YAC12-L) and circular (YAC12-C) configurations. **D.** In selective media (SC–His), SynIII strains carrying YAC12–eNeochrome III.v2, in either linear or circular form, display growth indistinguishable from the SynIII+pRS413-HIS3 as a control strain. Error bars indicate ± SEM (n = 3 biological replicates). **E.** Spot assays demonstrate a consistent healthy phenotype in SynIII strains harboring eNeochrome III.v3 or III.v4 on YAC12 with orthogonal REs, in both linear and circular forms. **F.** Corresponding growth curves show that SynIII strains expressing eNeochrome III.v3 or III.v4, in both linear and circular forms grow to levels comparable to the SynIII+pRS413-HIS3 control in selective media. Error bars indicate ± SEM (n = 3 biological replicates).

### 7. Comprehensive Phenotypic Analysis of Synthetic Chromosome III Variants Under 14 Different Conditions

We conducted a comprehensive phenotypic analysis of SynIII variants carrying different versions of the neo-essential synthetic chromosome III, including both circular and linear configurations. Growth and viability were assessed under 14 environmental and chemical stress conditions designed to evaluate cellular fitness and stress response. Spot assays were performed on YPD+G418 or SC-His, depending on the auxotrophic marker present on the essential neochromosomes. For comparison, we used SynIII+pRS-eNeochrome III.V1 S.c. after one generation (1g), which we designated as SynIII+pRS-eNeochrome III.V1. S.c. (1g), and after 100 generations (100g), which we designated as SynIII+pRS-eNeochrome III.V1. S.c. (100g). Assays were conducted at 25 °C, 30 °C, and 37 °C to evaluate temperature sensitivity. Stress conditions included camptothecin, 6-azauracil, benomyl, hydroxyurea, methyl methanesulfonate (MMS), cycloheximide, sorbitol, hydrogen peroxide, acidic stress (pH 4.0), alkaline stress (pH 9.0), and growth on non-fermentable carbon sources (YPEG; 2% glycerol + 2% ethanol). For cycloheximide and oxidative stress assays, cells were pretreated for 2 h before plating. The functional relevance of each condition is summarized in Table 1 (Dejean et al., 2000; Exinger & Lacroute, 1992; Hohmann, 2002; Kane, 2006; Lundin et al., 2005; Morano et al., 2012; Pommier, 2006; Schneider-Poetsch et al., 2010). SynIII+pRS-eNeochrome III.V1. *S. c.* (100g) generally retained phenotypic characteristics comparable to control strains (Fig. 8A). However, the SynIII+pRS-eNeochrome III.V1. *S.c.* (1g) variant exhibited growth defects under most tested conditions. Unexpectedly, this strain showed growth comparable to the control in the presence of camptothecin, a DNA topoisomerase I inhibitor widely studied for its anticancer activity [60, 61]. A similar behaviour was observed when 6-azauracil was added to the medium. 6-Azauracil reduces intracellular GTP levels and interferes with transcriptional elongation in yeast (Exinger & Lacroute, 1992; Tansey, 2006). The mechanistic basis for these resistance phenotypes remains unclear and will require further transcriptomic and proteomic investigation. In contrast, both SynIII+pRS-eNeochrome III.V1. *S.c.* (1g) and SynIII+pRS-eNeochrome III.V1. *S.c.* (100g) failed to grow on non-fermentable carbon sources. Nanopore sequencing revealed the complete loss of mitochondrial DNA in these strains following integration of the pRS-eNeochrome III. V1. *S.c.* construct, consistent with the observed respiratory deficiency. Most strains harboring essential neochromosomes on the YAC12 vector exhibited near wild-type growth under most conditions, in both the linear (L) form, YAC12.L-eNeochrome III.V2 *S.c*, and the circular (C) form, YAC12.C-eNeochrome III.V2 *S.c* (Fig. 9B). A pronounced growth defect was observed in SynIII+YAC12.L-eNeochrome III.V2 *S.c* and a moderate defect in SynIII+YAC12.L-eNeochrome III.V4 *S.e* when exposed to camptothecin, whereas SynIII+YAC12.C-eNeochrome III.V3 S.p displayed a strong camptothecin-resistant phenotype similar to WT strain. Exposure to MMS resulted in increased DNA damage sensitivity in SynIII+pRS413 and SynIII+YAC12.C-eNeochrome III.V2 *S. c*, whereas the strain harboring the linear chromosome SynIII+YAC12.L-eNeochrome III.V2 *S. c* exhibited no detectable effect. MMS is a classical alkylating agent widely used to study DNA repair pathways [62]. In contrast, all other SynIII strains harboring different versions showed no significant difference compared to the control strain WT BY4742 and performed slightly better than SynIII + pRS-HIS3. Under alkaline stress (pH 9.0), growth was reduced for all strains, including controls. However, strains harboring essential neo-chromosomes constructed with regulatory elements derived from *S. cerevisiae* exhibited a more pronounced growth defect compared with other variants. The molecular basis for this differential response remains unclear and warrants further investigation through comprehensive omics analyses.

**Figure 8.**
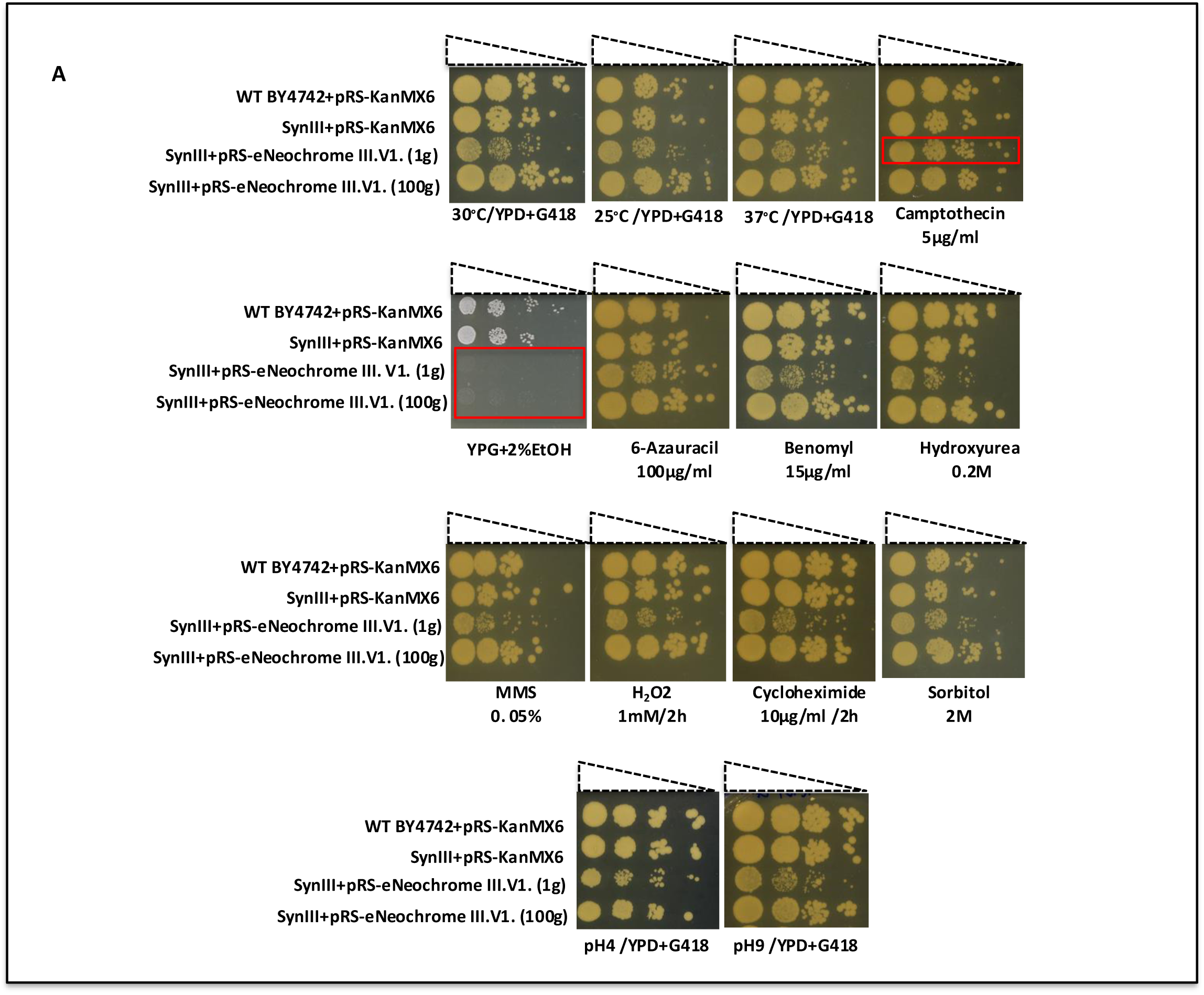
Comprehensive phenotypic analysis of SynIII strains harboring different versions of eNeochrome III (V1–V4) under diverse stress conditions. **A.** Comprehensive phenotypic analysis of SynIII strains harboring eNeochrome III.v1. Ten-fold serial dilutions were prepared from overnight cultures of SynIII strains carrying the neo-essential chromosome III, starting from an OD₆₀₀ of 0.1. Strains were identified by their corresponding strain IDs listed on the left. Spot assays were performed for SynIII strains harboring eNeochrome III.v1 on a pRS vector. Control strains carrying the empty pRS-KanMX6 vector (WT BY4742 + pRS-KanMX6 and SynIII + pRS-KanMX6) were included for comparison. Spot tests were conducted on selective YPD + G418 media at 25°C, 30°C, and 37°C, as well as under multiple stress conditions, including SM + camptothecin (5µg/ml), YPEG (2% glycerol + 2% ethanol), SM + 6-azauracil (100µg/ml), SM + benomyl (5µg/ml), SM + hydroxyurea (0.2M), YPD + MMS (0. 05%), SM + cycloheximide (10 μg/mL, 2 h pre-treatment), and YPD + H₂O₂ (1 mM, 2 h pre-treatment). Additional assays were performed on SM supplemented with sorbitol (2M), low pH (pH 4.0), and high pH (pH 9.0). The first-generation SynIII + pRS–eNeochrome III.v1 strain (1g), where g1 denotes the first generation of the constructed strain, exhibited a pronounced growth defect under most tested conditions. Notably, growth fitness was restored after prolonged passaging, as observed in the 100-generation strain (100g). Unexpectedly, the SynIII + pRS–eNeochrome III.v1-strain (1g) displayed similar growth to controls when exposed to YPD + G418 supplemented with camptothecin or 6-azauracil. Both SynIII + pRS–eNeochrome III.v1 strains (1g and 100g) failed to grow on media containing non-fermentable carbon sources as the sole energy source. Nanopore sequencing revealed loss of mitochondrial DNA in these strains following introduction of pRS–eNeochrome III.v1 into the semi-synthetic SynIII background. **B.** Comprehensive phenotypic analysis of SynIII strains harboring eNeochrome III.v2 *S.c*, III.v3 *S.p*, and III.v4 *S.e* engineered with native or orthogonal recombination elements (REs) derived from *S. cerevisiae*, *S. paradoxus*, and *S. eubayanus* on the YAC12 vector. Control strains carrying the empty vector with *HIS3*-marker (SynIII+pRS413 and WT BY4742+pRS413) were included for comparison. Ten-fold serial dilutions were prepared from overnight cultures of SynIII strains carrying different versions of the neo-essential chromosome III, starting from an OD₆₀₀ of 0.1. Strains were identified by their corresponding strain IDs listed on the left. Spot assays were performed on SC–His selective media and imaged after three days of incubation. Most SynIII strains harboring essential neochromosomes on YAC12 exhibited robust growth across tested conditions. However, a severe growth defect was observed in SynIII+YAC12.C– eNeochrome III.v2 (*S. cerevisiae*) and a moderate defect in SynIII+YAC12.L–eNeochrome III.v4 (*S. eubayanus*) upon camptothecin treatment. In contrast, SynIII+YAC12.C–eNeochrome III.v3 (*S. paradoxus*) displayed pronounced resistance to camptothecin. Both SynIII+pRS413 and SynIII+YAC12.C– eNeochrome III.v2 (*S. cerevisiae*) exhibited sensitivity to MMS, whereas SynIII+YAC12.L–eNeochrome III.v2 (*S. cerevisiae*) and SynIII+YAC12.C–eNeochrome III.v3 (*S. paradoxus*) showed resistance comparable to WT BY4742 + pRS413. Notably, strains harboring essential neo chromosomes with recombination elements derived from *S. cerevisiae* exhibited marked growth impairment at high pH (pH 9.0) compared with SynIII strains carrying refactored recombination elements. Although the underlying mechanism remains unclear, we anticipate that future transcriptomic and proteomic analyses will provide mechanistic insight. (move to the text)

**Figure 9.**
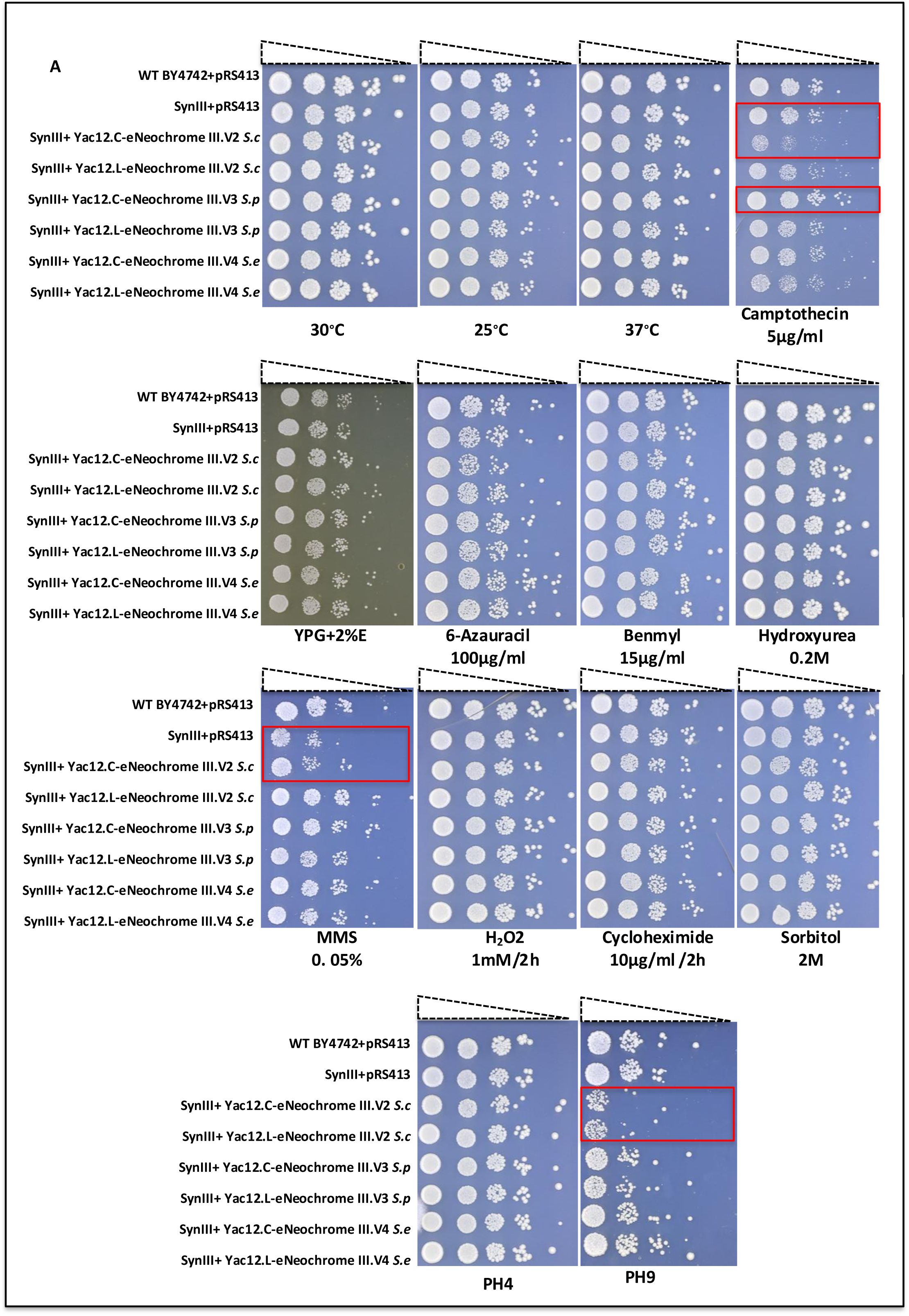
Stability of eNeochrome III variants (v1–v4) over 100 generations in linear and circular configurations. **A.** Schematic overview of the experimental workflow used to assess chromosomal stability of eNeochrome III variants. SynIII strains harboring different versions of eNeochrome III on either pRS or YAC vectors were propagated for 100 generations in the appropriate selective liquid media. After 100 generations, WT PCRTag analysis of chromosome III essential genes was performed on five biological replicates from SynIII strains carrying eNeochrome III variants (v1, v2, v3, and v4) in both linear and circular forms. This analysis confirmed the presence of all 14 essential genes from chromosome III engineered with native or orthogonal recombination elements, demonstrating high stability of all eNeochrome III variants under prolonged selective pressure. **B.** Phenotypic analysis shows complete recovery of the growth defect in SynIII + pRS–eNeochrome III.v1 *S. c* after 100 generations. **C.** Growth curve analysis (right panel) illustrates recovery of the growth defect in SynIII+pRS–eNeochrome III.v1 *S. c,* following 100 generations. **D.** Copy number variation analysis of an essential gene reveals that the copy number of *PGK1* is reduced from four to two after 100 generations in the SynIII + pRS–eNeochrome III.v1 S. c strain, buffering the impact on essential gene dosage and contributing to restoration of the phenotype.**E.** Phenotypic analysis demonstrates a consistent healthy phenotype for SynIII strains harboring YAC12-based eNeochrome III.v2 *S. c*, eNeochrome III.v3 *S. p*, and eNeochrome III.v4 *S. e* in both linear and circular configurations, comparable to the SynIII strain carrying the empty pRS413 vector.

**Table 1.** Drugs and stress conditions used for functional assays.

| Drug / Condition | Function / Cellular process tested | Reference |
| --- | --- | --- |
| Camptothecin | DNA topoisomerase I inhibitor; induces apoptosis and DNA damage by stabilizing Topo I–DNA cleavage complexes | (Pommier, 2006) |
| 6-Azauracil (6-AU) | Transcription elongation inhibitor; used to detect defects in RNA polymerase II transcriptional elongation | (Exinger & Lacroute, 1992) |
| Methyl methanesulfonate (MMS) | DNA alkylating agent; used to test DNA repair capacity | (Lundin et al., 2005) |
| Benomyl | Microtubule destabilizing agent; used to assess cytoskeletal integrity and chromosome segregation | Fiechter et al., 2008 |
| Sorbitol | Osmotic stress inducer; tests cell wall integrity and osmoregulation | (Hohmann, 2002) |
| YPEG medium | Non-fermentable carbon source medium; used to test mitochondrial respiration defects | (Dejean et al., 2000) |
| pH 4.0 / pH 9.0 (HCl/NaOH) | Acidic and alkaline stress; used to assess vacuolar function and pH homeostasis | (Kane, 2006) |
| Hydrogen peroxide (H<sub>2</sub>O<sub>2</sub>) | Reactive oxygen species; used to test oxidative stress response | (Morano et al., 2012) |
| Cycloheximide | Translation inhibitor; used to assess protein synthesis and ribosomal function | (Schneider-Poetsch et al., 2010) |

### 8. Chromosomal stability assay of circular and linear eNeochrome variants over 100 generations in SynIII

We evaluated the chromosomal stability of circular and linear versions of eNeochromeIII (eNeochrome V1–V4) over 100 generations under selective pressure. Overnight cultures of haploid SynIII strains harboring different eNeochromeIII variants were divided into two experimental groups. The first group included strains containing eNeochrome with native regulatory elements (RE) derived from *S.c.* in circular or linear form on pRS or YAC12 vectors: SynIII+pRS-eNeochromeIII V1. *S.c.*, SynIII+YAC12-C-eNeochrome.V2 *S.c.*, and SynIII+YAC12-L-eNeochrome.V2 *S.c*. The second group consisted of strains carrying eNeochrome with orthogonal RE in circular or linear configurations: SynIII+YAC12-C-eNeochrome III.V3 *S.p.*, SynIII+YAC12-L-eNeochromeIII.V3 *S.p*, SynIII+YAC12-C-eNeochromeIII.V4 *S.e.*, and SynIII+YAC12-L-eNeochromeIII.V4 *S.e*. Five biological replicates were used for each strain containing eNeochrome and were inoculated into 50 mL of selective medium (SC-His or YPD+G418) at a starting OD_600_ of 0.01. Every 24 h, cultures were back-diluted to OD_600_ 0.01. Glycerol stocks were prepared daily over a 10-day time course, corresponding to ∼10 generations per day and approximately 100 generations total [63]. On day 10, dilution series were plated on selective solid media. Five randomly selected colonies per strain were analyzed as biological replicates for genomic DNA extraction and WT PCRTag analysis. Under selective pressure, all circular and linear eNeochromeIII variants on both pRS and YAC vectors exhibited high stability over 100 generations (Fig. 10A). All 14 essential genes were retained with no detectable loss. Notably, SynIII+pRS-eNeochromeIII V1. *S.c.* displayed an initial growth defect that was fully rescued after 100 generations of adaptive evolution (Fig. 10B). Growth recovery of the evolved strain SynIII+pRS-eNeochromeIII V1. *S.c.* (100g) was confirmed in liquid culture assays (Fig. 10C). Whole-genome sequencing was performed on SynIII, SynIII+pRS-eNeochromeIII V1. *S.c.* (1g), and SynIII+pRS-eNeochromeIII V1. *S.c.* (100g) to identify mutations associated with growth recovery. Although multiple point mutations were detected in both strains compared to the control SynIII strain, none could be directly linked to the rescued phenotype (data not shown). We therefore performed copy number variation (CNV) analysis of the essential gene *PGK1* located on chromosome III. In haploid SynIII, *PGK1* is present as a single copy. CNV analysis revealed two copies of *PGK1* in SynIII+YAC12-C-eNeochromeIII.V2 *S.c.*, consistent with maintenance of the parental phenotype, as YAC-based constructs are typically maintained at low copy number [58]. In contrast, SynIII+pRS-eNeochromeIII. V1 *S.c.* (1g) carried approximately five copies of *PGK1*, reflecting elevated pRS vector copy number. After 100 generations, the evolved strain contained only two copies of *PGK1*. This reduction indicates adaptive evolution toward a tolerable essential gene dosage state, likely buffering the essential gene imbalance caused by the introduction of multiple pRS-borne eNeochromeIII copies into SynIII (Fig. 10D). This adaptation enabled restoration of growth in both rich and selective media, although the strain had completely lost mitochondrial DNA and was confirmed via nanopore sequencing. Importantly, SynIII strains harboring various versions of eNeochromeIII V2, V3 and V4 with native or orthogonal RE on the Yac12 vector consistently displayed robust growth from generation 1 through generation 100 (Fig. 10E). These strains maintained a healthy phenotype comparable to the SynIII + pRS413 control and exhibited slightly improved growth on MMs. We also confirmed by flow cytometry (FACS) that the strain harboring SynIII + pRS-eNeochromeIII V1. *S. c.* remained haploid at both the first and 100th generations. Similarly, the strain carrying eNeochromeIII V2 on YAC12 also remained haploid (Fig. 1S).

**Figure 10.**
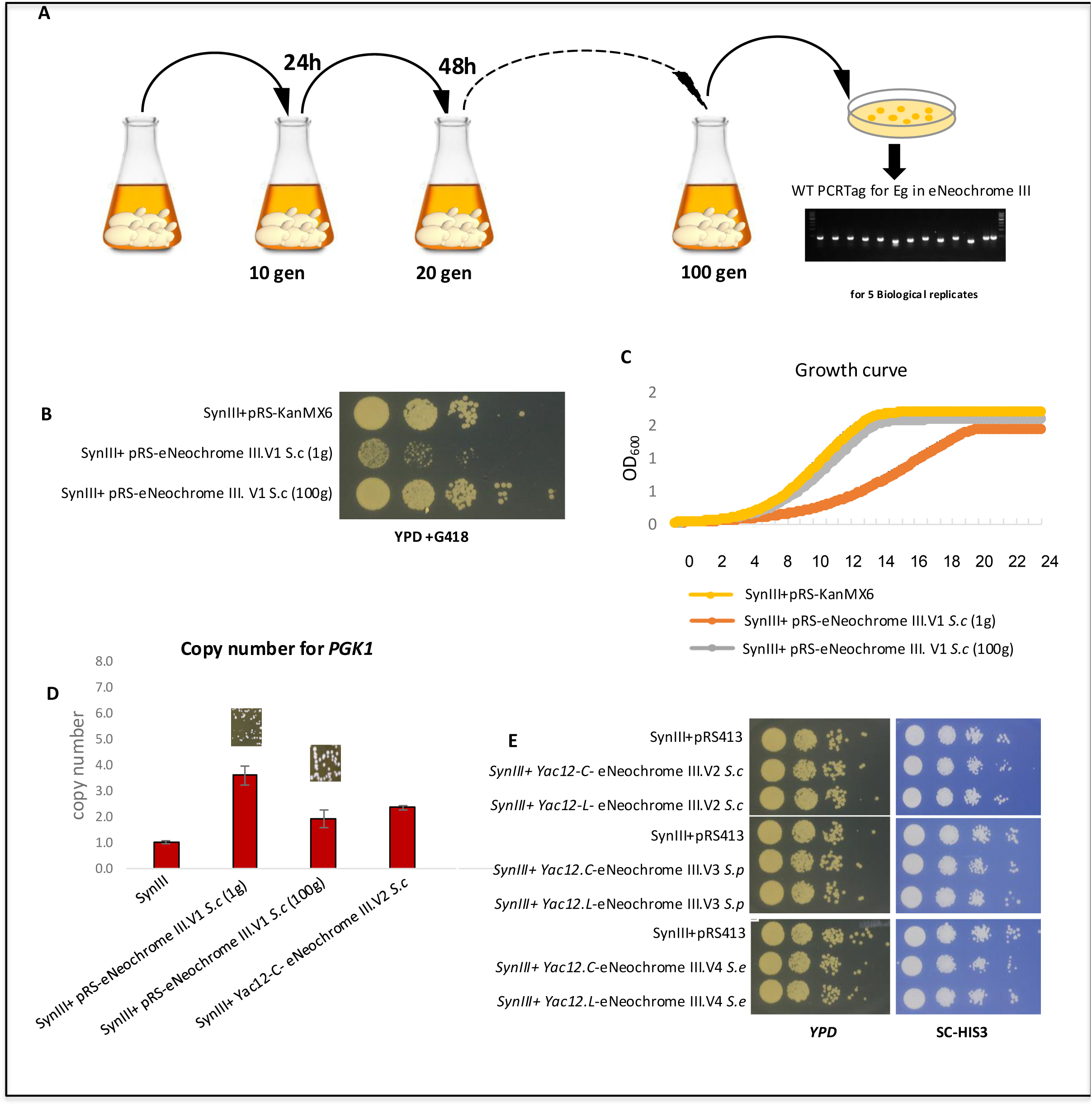
Workflow to minimise the synthetic genome by SCRaMbLE. **A.** Diagram of the SCRaMbLE reporter ERICA (Elementary Random Integration Cassette). The ERICA cassette contains a URA3 expression cassette (promoter–ORF–terminator) flanked by 34 bp loxPsym sites. These loxPsym sites enable random integration into any compatible loxPsym site within the synthetic chromosome via homologous recombination, eliminating the need for locus-specific oligo design. **B**. Schematic overview of the workflow used to minimize synthetic chromosome size. SynIII+pRS-eNeo.V1 *S. cerevisiae* serves as the starter strain. ERICA is randomly integrated into loxPsym sites on synthetic chromosome III to generate a SynIII strain harboring eNeoChromIII.V1, which is used as a pooled population. SCRaMbLE is induced for 3, 5, 7, 24, or 72 hours using Cre recombinase driven by either the daughter-specific promoter pSCW11 or the strong constitutive *TDH3* promoter. Post-SCRaMbLE mutants that have lost the reporter are positively selected on 5-FOA medium. PCRTag analysis of genomic DNA is used to estimate gene deletions, and nanopore sequencing resolves structural variation, including deletions, duplications, inversions, and insertions. These data reveal deletion patterns and identify SCRaMbLE hot and cold spots. Mutants carrying the highest deletion burden are subjected to sequential rounds of SCRaMbLE. **C.** Analysing post-SCRaMbLEd mutants harboring minimal genomes using nanopore technology

### 9. Development of the SCRaMbLE reporter ERICA (Elementary Random Integration Cassette)

A major challenge in isolating SCRaMbLE mutants is that only a fraction of cells in an induced population undergoes Cre-mediated rearrangement [14, 39]. To initially address this limitation, we integrated a *URA3* cassette expression module (promoter, ORF, and terminator) as a SCRaMbLE reporter into SynIII between the ambiguous ORF *YCR006C* and the uncharacterized ORF *YCR007C* (∼102–108 kb on chromosome III). Integration of the *URA3* cassette did not affect strain fitness. We performed SCRaMbLE following 24 h induction of Cre recombinase under the control of the daughter-specific promoter pSCW11. SynIII + pRS-eNeochrome III.V1 S.c (1g) exhibited limited SCRaMbLEd rearrangements (Fig. X), with SynIII used as the control strain. Although the introduction of pRS-eNeochrome III. V1 *S.c* resulted in a measurable growth defect. SCRaMbLE induction was successful in all strains, as evidenced by the complete loss of the *URA3* reporter, and post-SCRaMbLE mutants that had lost the *URA3* reporter were positively selected on 5-FOA medium confirming e<icient screening. A similar *URA3*-based reporter strategy has been described previously [14]. Analysis of loxPsym-mediated deletions revealed total genome reductions of 6.45 kb, 9.55 kb, and 6.47 kb across three independent SCRaMbLEd strains **(Fig. S1A, B).** These deletions predominantly a<ected nonessential open reading frames, including YCR007C, YCR011C, YCR005C, and YCR006C, as well as the dubious ORF YCR013C. Notably, two strains (SCRaMbLEd1 and SCRaMbLEd2) carried deletions encompassing **YCR012W *(PGK1*)**, an essential glycolytic gene. Despite loss of this essential locus, both strains remained viable and maintained near-normal growth, indicating e<ective functional complementation. Such deletions were not observed prior to the introduction of pRS-eNeochrome III.V1 S.c, demonstrating that the neochromosome expands the accessible landscape of genomic rearrangements. Consistently, genes within loxPsym units 6, 44, 48, 43, 90, and 99 were dispensable under the tested conditions and could be removed without detectable fitness consequences **(Fig. S1C).** However, the use of a locus-specific URA3 SCRaMbLE reporter introduces important limitations. Selection enriches only for mutants that have lost the *URA3* cassette at the targeted locus, while rearrangements occurring elsewhere in the synthetic chromosome remain undetected. In addition, the extensive structural variation generated after each SCRaMbLE cycle often necessitates long-read sequencing to map rearrangements and redesign new integration oligonucleotides for subsequent reporter insertion of integration constructs, making iterative screening labor-intensive. To overcome these limitations, we developed a versatile SCRaMbLE reporter system termed **ERICA** (Elementary Random Integration Cassette). ERICA integrates randomly between loxPsym sites via homologous recombination, eliminating the need for locus-specific redesign in successive SCRaMbLE rounds. This enables population-level screening, where a pool of strains carries the reporter at diverse genomic positions, thereby increasing the range of detectable rearrangements. The ERICA cassette consists of a *URA3* expression module (promoter, ORF, and terminator) flanked by 34-bp loxPsym sequences, allowing e<icient integration at any loxPsym site. The cassette can be generated by PCR and directly transformed into SynIII + pRS-eNeochrome III.V1 S.c (100g) (Fig. 11A). The resulting ERICA-integrated strain pool was subjected to positive selection on 5-FOA plates, enabling recovery of diverse SCRaMbLEd candidates that had lost ERICA from di<erent chromosomal loci. After recovery of a healthier phenotype after 100 generations, SynIII + pRS-eNeochrome III.V1 S.c (100g) was selected for further SCRaMbLE experiments, as it displayed a more robust phenotype compared with the unevolved 1-generation strain. SCRaMbLE was therefore performed using this evolved background. The ERICA cassette was amplified by PCR and transformed into SynIII + pRS-eNeochrome III.V1 S.c (100g). Cre recombinase was introduced under the control of either the daughter-specific promoter pSCW11 or the constitutive *TDH3* promoter in separate transformations. Induction was carried out for 3, 5, 7, 24, and 72 h, followed by positive selection of post-SCRaMbLE mutants on 5-FOA medium to isolate strains that had lost the *URA3* SCRaMbLE reporter. ERICA was subsequently applied for genome minimisation via SCRaMbLE. Using SynIII + pRS-eNeochrome III.V1 S.c (100g) as a pooled population, we identified 18 out of 19 bona fide SCRaMbLEd strains exhibiting successful reporter loss and genomic rearrangements. Importantly, ERICA demonstrates several design advantages over conventional locus-specific *URA3* reporters. The cassette can be readily generated as a PCR product and efficiently transformed without additional cloning steps. ERICA enables random integration via homologous recombination between loxPsym sites, eliminating the need for locus-specific targeting. This allows the use of pooled populations, substantially increasing the number and diversity of SCRaMbLEd variants that can be screened. In contrast, traditional *URA3* reporter systems integrated at a fixed locus restrict detection to a limited subset of rearrangements and significantly reduce screening throughput.

**Figure 11.**
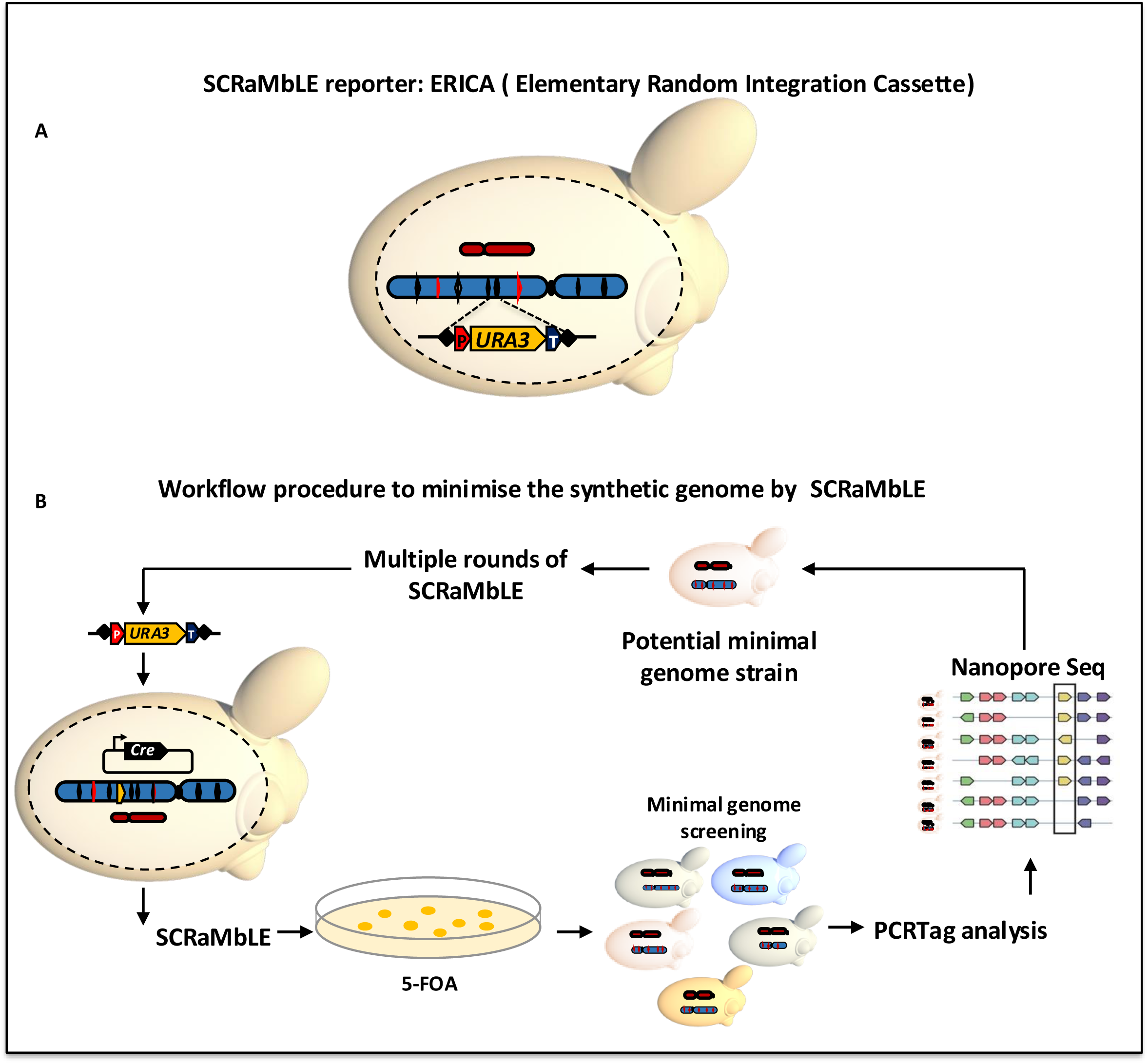
SCRaMbLE-mediated minimization of synthetic chromosome III in the synIII (Sc2.0) strain. **A.** Schematic representation of synthetic chromosome III showing essential genes in their native orientations (red) and regions containing nonessential genes (blue). The top track depicts the control synIII strain; the centromere is indicated by a black line near the left arm. Red arrows indicate essential genes, while blue arrows indicate nonessential genes; arrow length is proportional to gene size. IDs of post-SCRaMbLE mutants are shown on the left, with SCRaMbLE-induced structural variants, including deletions, duplications, and insertions, displayed relative to the control synIII chromosome. **B.** The figure shows the size distribution of deletions in post-SCRaMbLE mutants generated by SCRaMbLE. **Table 2**. Summary of structural variation and essential gene loss in post-SCRaMbLE mutants.

**Table 2.** Comprehensive Structural rearrangements and gene deletions in post-SCRaMbLEd mutants.

| Mutant ID | Essential gene deleted | Structural rearrangements | Total deletion (bp) | Deleted genes | Functional annotation |
| --- | --- | --- | --- | --- | --- |
| RSY521 | None | Insertion | – | – | – |
| RSY551 | None | None | – | – | – |
| RSY552 | None | 3 Inv (2964, 82029, 21231) | – | – | – |
| RSY575 | None | None | – | – | – |
| RSY577 | 3 essential genes: **YCR052W**, **YCR054C** and **YCR072C** | 3 Inv (67488, 3242, 33483); Dup (39963); Del (39930) | 39,930 | YCR047W-A, YCR047C, YCR048W, YCR049C, YCR050C, YCR051W, **YCR052W**, YCR053W, **YCR054C**, YCR059C, YCR060W, YCR061W, YCR063W, YCR064C, YCR065W, YCR066W, YCR067C, YCR068W, YCR069W, YCR071C, **YCR072C**, YCR073C, YCR073W-A, YCR075C, YCR075W-A, YCR076C | Predominantly uncharacterized ORFs; some linked to metabolism, transcription, and stress response |
| RSY578 | None | Del (219); Inv (750) | 219 | – | – |
| RSY579 | None | Inv (67490); Del (6000) | 6,000 | YCR005C, YCR006C, YCR007C | Unknown / poorly characterized proteins |
| RSY642 | None | 2 Del (33, 220); Inv (397) | 253 | – | – |
| RSY649 | None | Del (3286); 4 Inv (2530, 11265, 315, 2486) | 3,286 | YCR007C | Unknown function |
| RSY650 | **YCL004W (essential)** | 2 Del (218, 21230); Dup (27316) | 21,448 | YCL008C, YCL007C, YCL005W-A, YCL005W, YCL004W, YCL002C, YCL001W, YCL001W-A, YCL001W-B, YCR001W, YCR002C, YCR003W, YCR004C | YCL004W (essential): required for cell viability<br>Others: ribosome biogenesis, transcription, metabolism, unknown ORFs |
| RSY651 | None | Del (4712); Dup (8875); Inv (1689) | 4,712 | YCL027W, YCL026C, YCL026C-A | RNA processing / unknown proteins |
| RSY652 | None | Del (86); Dup (8842, 15080); Inv (316, 11496) | 86 | – | – |
| RSY671 | None | Inv (315) | – | – | – |
| RSY673 | None | Inv (315) | – | – | – |
| RSY676 | None | Inv (315) | – | – | – |
| RSY678 | None | Inv (315) | – | – | – |
| RSY680 | **PDI1 (YCL031C) (essential)** | 5 Del (86, 1480, 349, 8086, 6947); 4 Inv (5016, 8630, 11079, 131366); Ins (117); Dup (15500) | 26,648 | **YCL031C**, YCL030C, YCR021C, YCR022C, YCR023C, YCR024C, YCR024C-A, YCR024C-B, YCR025C, YCR084C, YCR085W, YCR086W, YCR087W, YCR087C-A, YCR088W | PDI1 (essential): ER protein folding (disulfide bond formation)<br>Others: protein processing, metabolism, transcription, unknown ORFs |
| RSY683 | None | Inv (315); InvDup (100); multiple insertions | – | – | – |
| RSY685 | None | Inv (315) | – | – | – |

### 10. Genome minimization via SCRaMbLE, validated using long-read nanopore sequencing

Chromosome III is the third smallest chromosome in *S. cerevisiae* and was the first eukaryotic chromosome to be sequenced (Oliver et al., 1992). In the Sc2.0 project, synthetic chromosome III (SynIII) is 272,871 bp, representing a 13.8% size reduction relative to the native chromosome. Sequence redesign occurs approximately every 500 bp and accounts for ∼2.5% total sequence modification, yet SynIII retains wild-type growth and fitness [13]. Structurally, chromosome III is a linear submetacentric chromosome with the centromere (*CEN3)* positioned toward the left arm, generating unequal arm lengths [64, 65]. SynIII contains 106 annotated genes, including 77 non-essential and 14 essential genes. The essential gene *SUP61* was removed from SynIII and temporarily relocated to the *HO* locus and will be reintegrated into the tRNA neochromosome [19]. The chromosome contains 98 loxPsym sites, of which 12 harbor essential genes. These essential loxPsym units, termed essential rafts, previously could not be deleted without lethality [13]. Units 8 and 73 each contain two essential genes, unit 69 spans ∼15 kb, and the centromere resides within unit 37. To screen genome minimization, 69 synthetic PCRTags were used, each corresponding to a gene within an individual loxPsym unit. PCRTag analysis detects deletions larger than 1–2 kb, while smaller rearrangements were resolved by nanopore sequencing. SCRaMbLE induction promotes Cre-mediated recombination between loxPsym sites positioned downstream of non-essential genes. Genome minimisation was performed using SCRaMbLE in SynIII + pRS-eNeochrome III.V1 *S.c* (100g). SCRaMbLE was induced for 3, 5, 7, 24, and 72 h using Cre recombinase driven by either the daughter-specific promoter pSCW11 or the constitutive promoter p*TDH3.* Mutants that lost the ERICA cassette were selected on 5-FOA medium, and gene deletions were assessed by PCRTag analysis. In the SynIII control strain lacking pRS-eNeochrome III.V1 *S.c*, essential loxPsym units remained undeletable, as confirmed by PCRTag. In contrast, in the presence of pRS-eNeochrome III.V1 *S.c*, deletions encompassing some essential gene regions were detected. Selected mutants were further validated by long-read nanopore sequencing to exclude PCR artefacts and to resolve structural variation, including deletions, duplications, inversions, and insertions **(Fig. 12A).** Across the mutant collection, diverse structural rearrangements were observed, reflecting the high combinatorial diversity generated by SCRaMbLE. Deletion sizes ranged from small events (86–253 bp) to large-scale genomic losses exceeding 39 kb. After a single SCRaMbLE round, deletions reached ∼40 kb in RSY577, ∼27 kb in RSY680, and ∼21.4 kb in RSY650 **(Fig. 12B**; **Table 2).** These deletions predominantly affected nonessential or poorly characterised open reading frames. For example, RSY579 and RSY649 exhibited deletions affecting YCR005C–YCR007C, while RSY651 involved the removal of loci within the YCL026–YCL027 region. Strikingly, several strains tolerated deletion of essential genes. RSY577 exhibited a 39.9 kb deletion encompassing three essential genes (**YCR052W, YCR054C, and YCR072C**), together with multiple inversions and a duplication event for the deleted locus. Similarly, RSY650 and RSY680 carried deletions affecting essential genes, including **YCL004W** and YCL031C (***PDI1)***, respectively. Despite loss of these essential loci, all strains remained viable, indicating robust functional complementation. In addition to deletions, complex rearrangements were frequently observed. RSY577 and RSY680 displayed combinations of inversions, duplications, insertions, and deletions, whereas other strains showed simpler architectures, such as single inversions (e.g., RSY671–RSY685), highlighting variability in genomic complexity across the population. Collectively, these results demonstrate that SCRaMbLE, coupled with the ERICA reporter system, enables extensive genome restructuring, including large-scale deletions and complex rearrangements. Notably, the tolerance to simultaneous deletion of multiple essential genes underscores the neochromosome’s capacity to buffer essential functions and expands the accessible landscape of genome minimisation.

**Figure 12.**
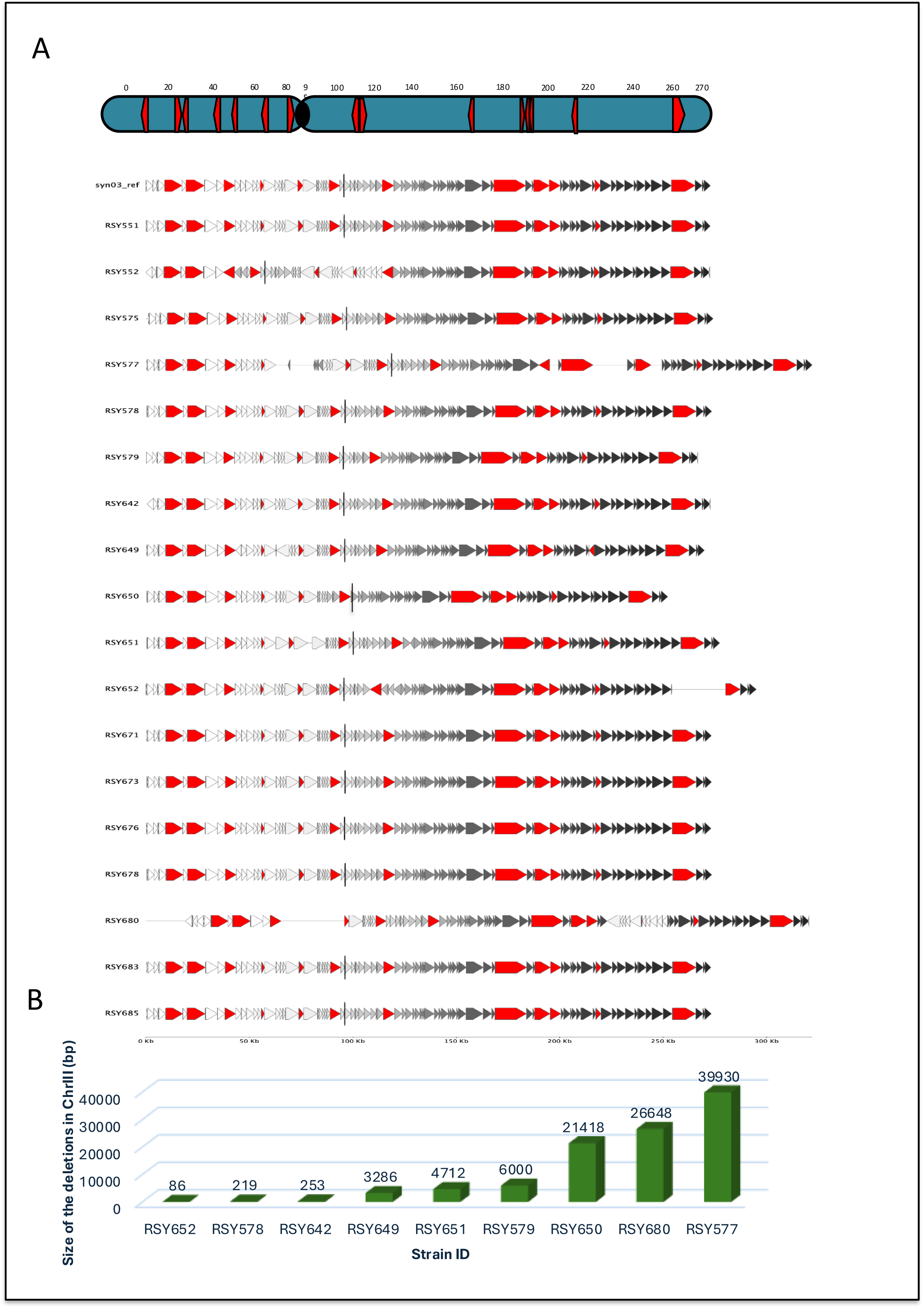
CREePY system demonstrates high deletion efficiency at small scale for removal of native *ADE2* and synthetic *FUS1* on chromosome III using marker-based and marker-less deletion strategies. **A.** Schematic representation of the deletion strategy used to evaluate deletion efficiency of the CREePY CRISPR/Cas9 system. The CRISPR/Cas9 vector harbors two gRNAs targeting the gene to be deleted and is co-transformed with a DNA repair template (1,000 bp) consisting of 500 bp left and right homology flanking regions (HFRs). Both constructs were transformed into synIII+pRS–eNeochrome III.v1 *S. c*, and transformants were selected on SC-URA medium for CREePY. **B**. Schematic representation of the marker-based deletion strategy used to evaluate deletion efficiency of the CREePY CRISPR/Cas9 system. The CRISPR/Cas9 vector harbors two gRNAs targeting the gene to be deleted and is co-transformed with a DNA repair template containing a deletion marker flanked by 500 bp left and right HFRs of the target gene. Both constructs were transformed into synIII+pRS–eNeochrome III.v1 *S. cerevisiae*, and transformants were selected on SC-URA-LEU medium for CREePY. **C.** Comparison of deletion efficiency for the native gene *ADE2* and the synthetic gene *FUS1* using either a *LEU2* deletion cassette or a markerless cassette with the CREePY system. Negative controls are shown on the right. Deletions were verified by confirmation PCR of both upstream and downstream integration sites. Deletion efficiency for both strategies and both genes was approximately 99%.

### 11. CREePY CRISPR/CasG supports near-complete small-scale deletion efficiency, but large chromosome III excisions remain constrained

To quantify the deletion efficiency of the CREePY CRISPR/Cas9 platform [66], we performed proof-of-concept excisions targeting both a native locus (*ADE2*) in chromosome XV and a synthetic locus (*FUS1*) in chromosome III in the synIII+pRS–eNeoChromIII.V1 *S. c* strain. Deletion of *ADE2* provides a visual phenotypic marker because loss of gene function produces pink colonies, allowing rapid enrichment of candidate transformants before molecular confirmation. Two repair strategies were compared: a markerless repair template containing 500 bp homology flanking regions (HFRs) and a marker-based deletion cassette carrying a selectable marker flanked by identical HFRs, 500pb **(Fig. 13A–B).** The CRISPR/Cas9 vector was multiplexed with two guide RNAs positioned on either side of the target locus and co-transformed with the appropriate repair template. Transformants were selected on SC-URA medium for markerless deletions and SC-URA-LEU medium for marker-based deletions. A total of 20 independent colonies were screened as biological replicates for each deletion. Successful excision was verified by PCR across both upstream and downstream integration junctions (Fig. 13C). Both strategies yielded exceptionally high deletion efficiencies (∼99–100%) for *ADE2* and *FUS1*, confirming that the CREePY system enables highly reliable removal of both native genes in the native chromosome and synthetic gene in synthetic chromosome. These efficiencies are consistent with prior reports of near-complete editing using multiplex CRISPR systems in yeast when homologous repair is optimised [66, 67]. Following validation of small-scale editing performance, we extended the approach to large chromosomal deletions spanning 20–90 kb on chromosome III (Fig. 14A). For each targeted region, the CRISPR/Cas9 vector encoded three guide RNAs located at the beginning, midpoint, and end of the deletion interval to promote efficient excision. The repair template consisted of a *HIS3* deletion cassette flanked by extended 1.5 kb HFRs to increase homologous recombination efficiency (**Fig. 14B**). Transformants were recovered at high frequency under SC-URA-HIS selection, while no colonies were observed in the negative control, SynIII + pRS-eNeochrome III.V1 S.c (100g), lacking both the CREePY vector and the DNA repair template. This control was used to assess the frequency of false-positive transformants generated during transformation, confirming correct selection stringency. More than 125 independent colonies were screened as biological replicates. However, two-stage screening revealed that all candidates represented false positives. While primary PCR was to confirm integration of the deletion cassette, secondary synIII PCRTag analysis demonstrated retention of the targeted chromosomal regions, indicating ectopic cassette integration rather than correct locus replacement. Deletion success declined sharply with increasing segment size and essential gene density (Table 3), consistent with known genome-wide constraints on large chromosomal excisions in yeast [35, 37]. During strain characterisation via nanopore sequencing, we further established that synIII+pRS–eNeoChromIII.V1 *S.c* exhibited a petite phenotype and had completely lost mitochondrial DNA. The strain was unable to grow on non-fermentable carbon sources, confirming exclusive dependence on fermentation. Respiratory-deficient petite strains exhibit altered energy metabolism and reduced tolerance to cellular stress, conditions known to exacerbate sensitivity to genome instability and large-scale genetic perturbation [68]. Loss of mitochondrial function, therefore, likely imposed an additional physiological constraint contributing to the failure to recover large chromosome III deletions. We also confirmed by flow cytometry (FACS) that SynIII +pRS– eNeoChromIII.V1 *S.c* remained haploid after transformation and deletion events **(Fig. 2S).** Together, these findings demonstrate that the CREePY platform is highly effective for precise small-scale genome editing in synthetic yeast chromosomes, but that extensive chromosomal minimisation remains strongly restricted probably due to the impact of synthetic lethal interactions. The inability to obtain large deletions likely reflects a combined effect of essential gene density, structural genome stability limits, and metabolic fragility associated with mitochondrial DNA loss. Future optimisation may require restoration of mitochondrial function through backcrossing with a respiration-competent strain, followed by sporulation to isolate haploid progeny retaining both the synthetic chromosome and functional mitochondria. In addition, staged deletion of smaller intervals (5–10 kb) may reduce lethal interactions and improve recovery of viable minimisation mutants.

**Figure 13.**
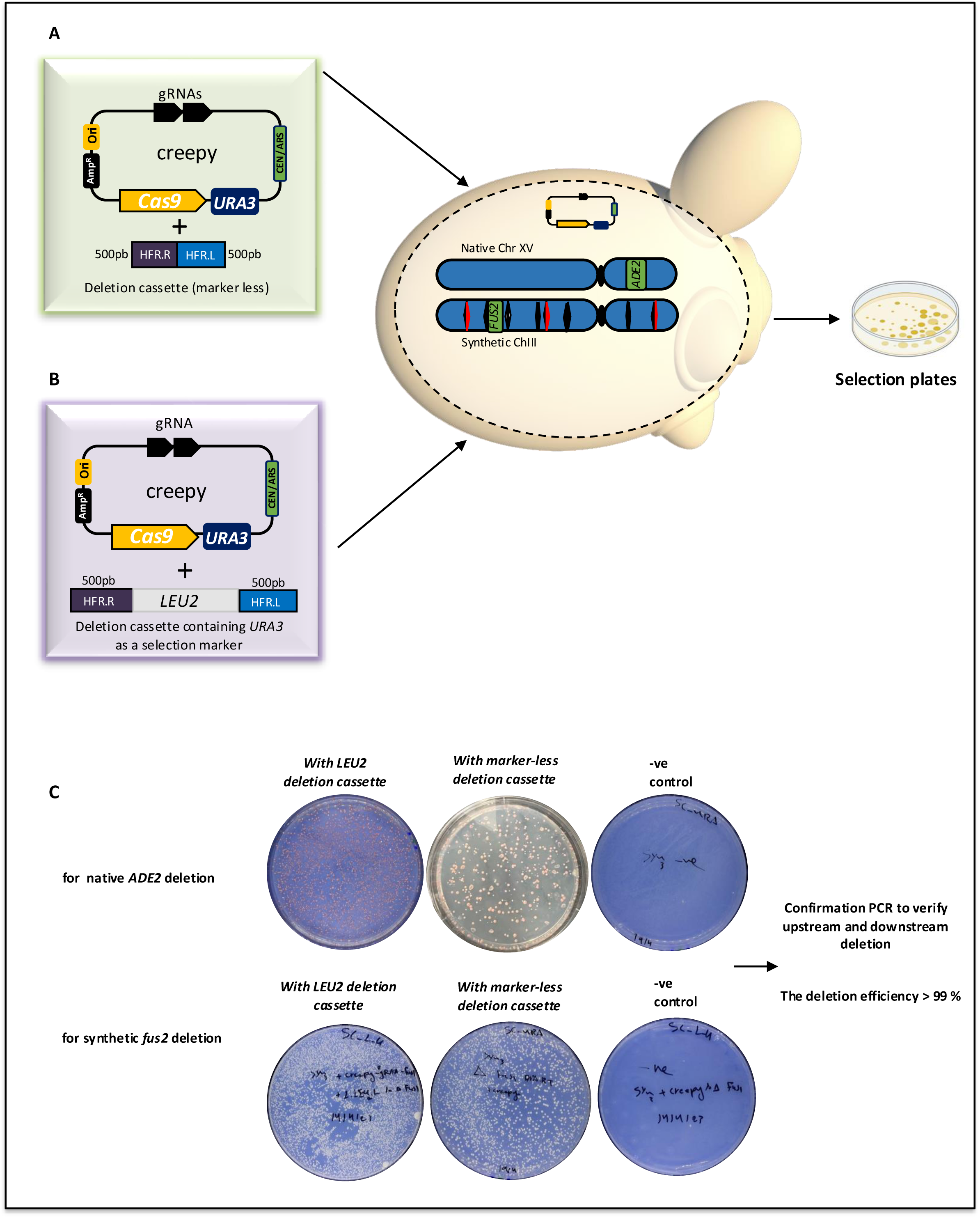
The figure shows the workflow used to delete large segments (20, 30, 40, 50, and 90 kb) of chromosome III. **A** Schematic of chromosome III highlighting essential genes and the region targeted for deletion. **B.** The CRISPR/Cas9 vector harbors three gRNAs targeting the locus to be deleted and is co-transformed with a DNA repair template containing a deletion marker (*HIS3*) flanked by long homology flanking region 1500 bp left and right (HFRs). Both constructs were co-transformed into synIII+pRS– eNeochrome III.v1 *S. c* (100g), and transformants were selected on SC-URA-HIS medium **C.** Colonies were initially selected from selective plates and subjected to primary screening by confirmation PCR to verify upstream and downstream integration sites, followed by secondary screening using synIII PCRTag analysis to confirm deletions by the absence of PCRTag products corresponding to the deleted regions. Table 1 summarizes deletion sizes, the number of essential genes within each region, and the number of mutants screened by confirmation PCR and synIII PCRTag analysis.

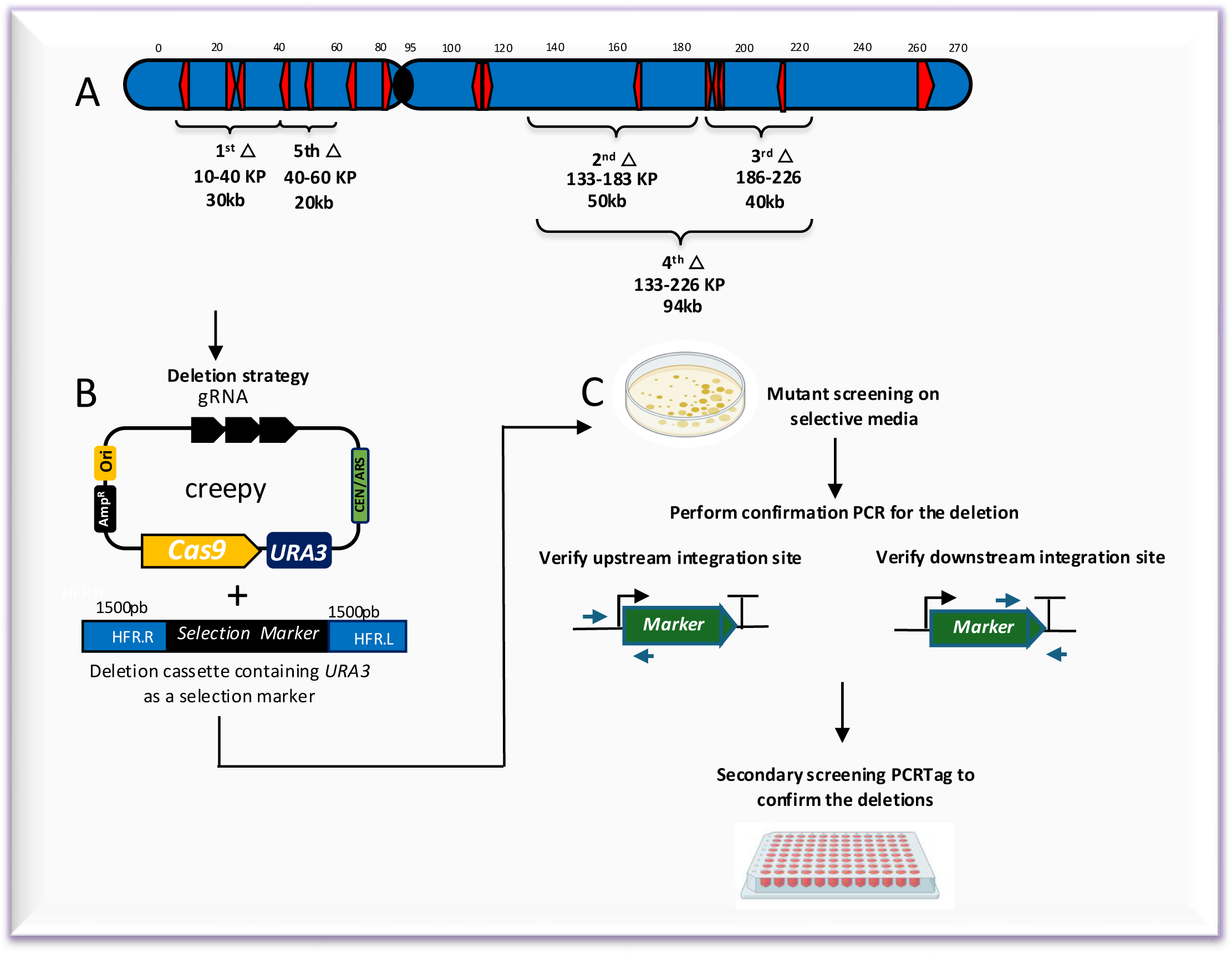

**Table 3.**
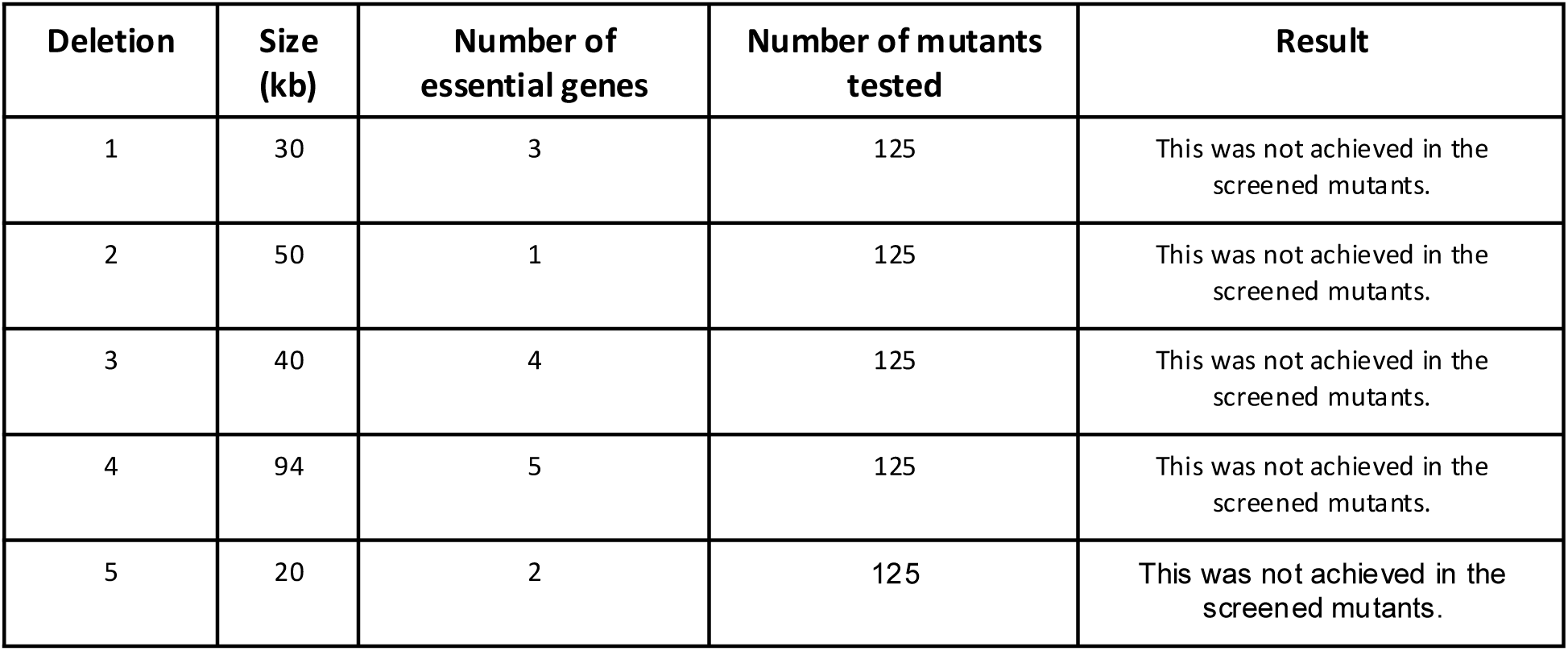
summarizes deletion sizes, the number of essential genes within each region, and the number of mutants screened.

| Deletion | Size (kb) | Number of essential genes | Number of mutants tested | Result |
| --- | --- | --- | --- | --- |
| 1 | 30 | 3 | 125 | This was not achieved in the screened mutants. |
| 2 | 50 | 1 | 125 | This was not achieved in the screened mutants. |
| 3 | 40 | 4 | 125 | This was not achieved in the screened mutants. |
| 4 | 94 | 5 | 125 | This was not achieved in the screened mutants. |
| 5 | 20 | 2 | 125 | This was not achieved in the screened mutants. |

### 12. Limitations of genome minimization via SCRaMbLE

Despite improving the deletion capacity of SCRaMbLE to remove essential genes in the presence of eNeoChromIII.V1 *S.c.,* extensive minimisation of synthetic chromosome III remains biologically constrained. SCRaMbLE is a powerful strategy for rapidly identifying dispensable genomic regions, enabling simultaneous interrogation of multiple loci and revealing functional dependencies that would be difficult to uncover through iterative gene-by-gene deletions. The analysis of SCRaMbLEd variants therefore provides valuable insights into genome organisation and essentiality. However, SCRaMbLE should be regarded primarily as a discovery tool rather than a direct method for constructing a minimal genome, as it generates complex and often unpredictable structural rearrangements alongside informative deletions. Large-scale gene removal increases the likelihood of synthetic-lethal interactions, where combinations of individually tolerated deletions become [5, 37]. Although yeast genomes exhibit substantial redundancy—approximately 80% of genes are individually non-essential in rich media—this buffering capacity has limits. Single deletions can sensitize cells to additional perturbations, environmental stress, or secondary gene loss [69, 70], such that simultaneous removal of multiple loci can overwhelm compensatory pathways and lead to loss of viability. More broadly, viability depends on network architecture, gene dosage balance, and chromosome integrity rather than single-gene essentiality alone. Essential functions are embedded within dense genetic interaction networks, and removal of extended chromosomal segments disrupts multiple interconnected pathways, sharply increasing lethality [36, 37]. Dosage sensitivity and haploinsufficiency further constrain tolerated deletions, while chromosomal context can amplify these effects. For example, SCRaMbLE-mediated rearrangements may reposition genes into repressive chromatin environments, leading to position-effect silencing that phenocopies essential gene loss [71]. In addition, chromosome stability imposes structural constraints on genome reduction. Maintenance of centromere, telomere, and replication functions is essential for viability [57], limiting the range of deletions that can be recovered. Physical properties of recombination also bias SCRaMbLE outcomes: Cre recombinase preferentially recombines loxPsym sites that are in close three-dimensional proximity within chromosomal interaction domains [72]. Similar distance-dependent effects have been observed in mammalian systems (Zheng et al., 2000), suggesting that genome topology influences which rearrangements are preferentially generated. Consequently, deletion landscapes are shaped not only by selection but also by spatial genome organization. From a practical perspective, SCRaMbLE generates a large diversity of rearrangements, but only a small fraction of the population undergoes recombination, creating an inherent screening bottleneck. Traditional locus-specific *URA3* reporters further restrict detection to rearrangements occurring at predefined sites, limiting the diversity of recoverable mutants. While the ERICA reporter alleviates some of these constraints by enabling random, locus-independent integration and pooled screening, distinguishing true SCRaMbLE-derived mutants from spontaneous *URA3* inactivation events remains challenging, and low-frequency false positives cannot be eliminated. Collectively, these biological, structural, and technical constraints explain why recovered mutants are enriched for small, viability-compatible rearrangements, whereas extensive chromosome III minimization remains difficult. Large deletions are likely generated at the DNA level but are selectively lost because only viable configurations can be recovered. As a result, SCRaMbLE-derived datasets represent a filtered landscape of genome architectures compatible with essential network integrity, underscoring both the power and the limitations of this approach for genome minimization [15, 17, 73].

## Discussion

Genome minimization aims to reduce genetic complexity while preserving cellular viability, thereby enabling the construction of simplified and predictable biological systems. Although approximately 80% of *Saccharomyces cerevisiae* genes are individually non-essential under rich growth conditions (Giaever et al., 2002; Winzeler et al., 1999), essentiality is highly context-dependent. Genetic interaction networks impose constraints such that combinations of individually tolerated deletions can become lethal, defining a fundamental boundary for genome reduction. A central advance of this study is the engineering of Neo-chromosome III variants that relocate essential genes under both native and orthogonal regulatory elements. Promoters derived from *S. paradoxus* and *S. eubayanus* supported expression comparable to their *S. cerevisiae* counterparts, demonstrating that essential gene function can be decoupled from native chromosomal context and regulatory architecture. This establishes a modular framework in which essential loci can be repositioned and reprogrammed without compromising viability, enabling programmable restructuring of synthetic chromosomes. SCRaMbLE provides a powerful platform for interrogating genome architecture at scale by enabling combinatorial deletions and rearrangements that uncover functional dependencies and viable genome configurations. Importantly, the introduction of eNeoChromIII.V1 *S.c* expands the deletion landscape by externalizing “essential rafts”—genomic regions in which essential and non-essential genes are interspersed between tow loxpSym units in the synthetic chromosome. In the absence of the eNeoChromIII.V1 *S.c*, these regions were refractory to deletion, as removal of such segments inevitably resulted in loss of essential gene function. In contrast, the presence of the eNeoChromIII.V1 *S.c* enabled deletion and restructuring of these regions, including large segments that were previously inaccessible. Gene-level analysis further refines the concept of essentiality within SynIII. Large deletions, such as those observed in RSY577, encompassed extensive gene clusters (YCR047–YCR076) that include both non-essential and essential genes. Notably, essential genes (**YCR052W, YCR054C, and YCR072C**) were deleted, and the whole segment was duplicated in chromosome III, yet tolerated, while YCL004W and YCL031C (*PDI1),* and YCR012W *(PGK1),* YCR013C were completely deleted in other strains. These findings demonstrate that essential gene function can be uncoupled from native chromosomal context and maintained through eNeoChromIII.V1 *S.c* -mediated complementation. In parallel, smaller recurrent deletions (e.g., YCR005C–YCR007C and YCL026–YCL027) define candidate dispensable modules that may be directly targeted for rational genome reduction. Together, these results suggest that genes can be categorized as truly dispensable, conditionally essential, or structurally relocatable depending on genomic context. Despite these advances, SCRaMbLE-derived genomes are structurally complex, frequently containing inversions, duplications, and insertions in addition to deletions. In some cases, deleted sequences may be partially retained or functionally compensated elsewhere in the genome, complicating interpretation of gene essentiality. These observations underscore that genome minimization is governed not only by gene identity but also by gene dosage, chromosomal context, and higher-order genetic interactions and the presence of synthetic lethal interaction. Consistent with previous studies (Wang et al., 2020), essential genes can tolerate rearrangement and relocation; however, accurate interpretation of these events requires high-resolution validation, as PCRTag-based approaches may fail to capture complex structural variation. Long-read sequencing is therefore essential for resolving genome architecture. Importantly, SCRaMbLE should be viewed as a discovery tool rather than a direct method for constructing minimal genomes. While it efficiently generates informative deletion landscapes, it also produces combinatorial rearrangements that can increase genome size or introduce unintended architectures. Nevertheless, this complexity represents a key strength, as it enables exploration of genetic interactions and epistatic relationships at a scale that is not tractable through manual engineering. Our results also highlight intrinsic biological constraints that limit large-scale genome reduction. Viability depends not only on single-gene essentiality but also on network architecture, gene dosage balance, and chromosome integrity. Removal of extended genomic regions disrupts multiple interacting pathways, increasing the likelihood of synthetic lethality even when individual genes are buffered (Costanzo et al., 2016; Giaever et al., 2002). Dosage sensitivity and haploinsufficiency further restrict tolerated deletions, while chromosomal context can amplify these effects through position-effect silencing (Gottschling et al., 1990). In addition, maintenance of centromere, telomere, and replication functions is essential for chromosome stability (Murray & Szostak, 1983), imposing structural constraints on genome minimization. These biological limitations are further underscored by our observations using the CREePY CRISPR/Cas9-based system. While CREePY enabled efficient excision of small genomic regions, attempts to generate large deletions frequently resulted in ectopic integration or compensatory genome rearrangements rather than stable removal of the targeted sequence. This highlights a key challenge: even when DNA cleavage is efficient, cellular repair mechanisms such as non-homologous end joining act to preserve genome integrity. Together, these findings indicate that the primary barrier to large-scale genome minimization is biological rather than technical. Consequently, SCRaMbLE-derived populations represent a filtered subset of viable genome configurations. Although large deletions are likely generated at the DNA level, only those compatible with essential network integrity, gene dosage balance, and chromosome stability are recovered (Annaluru et al., 2014; Richardson et al., 2017; Shen et al., 2016). This selection bias explains why extensive chromosome III minimization remains difficult despite high recombination efficiency. Looking forward, this work establishes a versatile platform for systematic exploration of genome architecture and essential gene function. Alternative versions of the essential neochromosome could be engineered to carry distinct regulatory configurations, gene sets, or synthetic modules, enabling controlled interrogation of essential gene buffering across diverse genomic contexts. In particular, different version of the neo-chromosomes could be introduced or exchanged into highly reduced SCRaMbLE-derived strains, for example via chromoduction-based approaches, to evaluate how different essential gene which refactored with orthogonal RE complements, influence cellular fitness, robustness, and adaptability. Such strategies would enable iterative refinement of genome minimization by decoupling essential function from chromosomal position and testing compatibility across multiple genomic backgrounds. More broadly, this framework provides a powerful foundation for functional dissection of synthetic chromosomes at unprecedented resolution. By combining SCRaMbLE-driven diversity with modular essential gene complementation, it becomes possible to systematically map genetic interactions, identify context-dependent essentiality, and probe the limits of genome plasticity. Beyond yeast, these principles may inform the design and functional characterization of artificial chromosomes in more complex eukaryotic systems. Together, this work positions synthetic chromosomes not only as engineering targets but also as experimental platforms for uncovering fundamental principles of genome organization, robustness, and evolution.

## Materials and Methods

### Strains, plasmids, synthetic gene fragments, and media

Laboratory strains used in this study included *Saccharomyces cerevisiae* wild-type BY4741 and the Sc2.0 semi-synthetic strain synIII harboring a synthetic chromosome. These strains were obtained from the Yizhi Cai laboratory (Manchester Institute of Biotechnology, University of Manchester, UK). The CREePY CRISPR/Cas9 expression system was derived from the Jef D. Boeke laboratory (Institute for Systems Genetics, NYU Grossman School of Medicine, USA) (Zhao et al., 2023). The pSCW11-cre-EBD and pTDH3-cre-EBD plasmids were introduced into synIII strains carrying eNeo-chromosome III to induce SCRaMbLE. Plasmids used for regulatory refactoring included pHCKan-P, pHCKan-O, pHCKan-T, pPOT vectors carrying URA3 or LEU2, pYFASS, and YAC12. Promoter activity was measured using the pRe2.8 YFP/mCherry dual-reporter system obtained from the Yizhi Cai laboratory (Manchester Institute of Biotechnology, University of Manchester, UK). Promoters and terminators from *S. paradoxus* and *S. eubayanus* were synthesized (Twist Bioscience, USA; Integrated DNA Technologies, USA) and cloned into standard vectors. Yeast strains were cultured using standard protocols. Non-selective growth was performed in YPD medium (10 g/L yeast extract, 20 g/L peptone, 20 g/L glucose). Selective growth was carried out in synthetic complete dextrose (SCD) medium supplemented with 2% glucose. G418 (Geneticin; Thermo Fisher Scientific, USA) or uracil dropout media were used where required. Media components were supplied by Fisher Scientific and Formedium (UK). Bacterial transformants were selected on LB medium supplemented with carbenicillin or kanamycin (Sigma-Aldrich, USA).

### Bacterial transformation

Chemically competent *E. coli* DH5α cells were used for plasmid propagation. Fifty microliters of cells were thawed on ice and mixed with 5 µL plasmid DNA (30–100 ng). After 30 min incubation on ice, cells were heat shocked at 42 °C for 45 s and returned to ice for 5 min. Cells were recovered in 950 µL SOC medium at 37 °C for 1 h with shaking and plated onto selective LB agar. For plasmids smaller than 10 kb, Mix & Go! DH5α competent cells (Zymo Research, USA) were used according to the manufacturer’s instructions.

### Restriction digest analysis

Plasmid DNA was digested using restriction enzymes from New England Biolabs (NEB, USA). Reaction conditions were selected using the NEB Double Digest Finder. Reactions contained up to 1 µg DNA, 1× NEBuffer, 0.5 µL of each enzyme, and nuclease-free water to 10 µL. Digestions were incubated for ≥1 h or overnight. Products were analyzed by 1% agarose gel electrophoresis.

### Assembly of YeastFAB parts

YeastFAB assembly followed previously described methods [50, 51]. PCR amplification used IDT primers and standard cycling conditions. Golden Gate assembly reactions included pHCKan-P/O/T parts and POT receiving vectors with BsmBI and T4 DNA ligase (Thermo Fisher Scientific). Verified constructs were screened by colony PCR, restriction digestion, and sequencing. Essential Neo-chromosome III fragments were assembled by homologous recombination following transformation into synIII. Correct assembly was confirmed by junction PCR and PCRTag analysis.

### Universal homologous region

The Universal Homologous Region (UHR) was designed to generate a 500 bp random DNA sequence. The UHR was incorporated into eNeochromosome III.v1 to facilitate the future assembly of essential genes from different chromosomes. Sequence specificity was assessed using BLAST (https://blast.ncbi.nlm.nih.gov/Blast.cgi) against the *Saccharomyces cerevisiae* genome, confirming no significant homology.The UHR was synthesised de novo from overlapping oligonucleotides using overlap-extension PCR, as described in the Build-A-Genome course.

### Yeast transformation

Yeast transformations were performed using the lithium acetate protocol [74]. Cells were grown to OD600 ≈ 1, washed, and transformed using LiOAc, PEG 3350, and herring sperm carrier DNA. Heat shock was performed at 42 °C for 30 min before plating onto selective media.

### DNA isolation

Genomic DNA was extracted using phenol–chloroform purification [15, 75]. Plasmid DNA was purified from bacterial cultures using QIAprep Spin Miniprep Kits (Qiagen, Germany).

### Growth curve analysis

Growth curves were recorded using a BioTek plate reader or a Tecan Infinite F200 microplate reader. Overnight cultures were diluted to OD_600_ ≈ 0.1 in 96-well plates with transpirable led and monitored over 24 h at 30°C. Experiments included three biological replicates with technical repeats.

### Phenotypic assays

Tenfold serial dilution spot assays were performed on selective and stress media. Chemical stressors included methyl methanesulfonate (MMS) **(0.05%)**, benomyl (15 µg/mL), camptothecin (5 µg/mL), hydroxyurea (0.2 M), sorbitol (2 M), 6-azauracil (100 µg/mL), hydrogen peroxide (1 mM), and cycloheximide (10 µg/mL) (all from Sigma-Aldrich, USA). For hydrogen peroxide and cycloheximide treatments, cells were exposed for 2 h before plating on selective media. Plates were incubated at 25 °C, 30 °C, or 37 °C for 3 days.

### Chromosomal stability assay

Chromosomal stability of circular and linear eNeochromosome III variants was evaluated over ∼100 generations under selective conditions. Haploid SynIII strains carrying different eNeochromosome III constructs were grown overnight at 30 °C and divided into experimental groups. Strains harboring eNeochromosomes with native regulatory elements derived from *S. cerevisiae* in either circular or linear form (on pRS or YAC12 vectors) were used. Five biological replicates per strain were inoculated into 50 mL of selective medium in 250 mL flasks (SC–His or YPD supplemented with G418) at an initial OD₆₀₀ of 0.01 and incubated at 30 °C with shaking (200 rpm). Cultures were propagated by daily back-dilution (1:1000) to OD₆₀₀ = 0.01 for 10 days (∼10 generations per day; ∼100 generations total) (Luo, Jiang, et al., 2021). Glycerol stocks were prepared daily throughout the experiment. On day 10, cultures were serially diluted, and 200 µL of the appropriate dilution was plated onto selective solid media and incubated at 30 °C. Five colonies per strain were randomly selected as biological replicates for genomic DNA extraction and WT PCRTag analysis.

### SCRaMbLE induction

SCRaMbLE was induced by adding β-estradiol (1 µM; Sigma-Aldrich, Germany) to cultures carrying the Cre-EBD plasmid under the control of either the daughter-specific pSCW11 promoter or the strong *TDH3* promoter. After 24 h of induction, cells were plated onto 5-FOA medium to select for post-SCRaMbLE mutants. Deletions were screened by PCRTag analysis.

### PCRTag analysis

To screen for candidate isolates, primers were designed to anneal ORF of the gene wt/synthetic to produce an amplicon of approximately 400 bp to 600 bp. Crude genomic DNA was generated by incubating yeast isolates at 95°C in 50 mL of 20 mM NaOH for 10 minutes or the MasterPure Yeast DNA Purification Kit (Lucigen). PCR Tag analysis was performed either by adding 1.8 µL of yeast lysate used as template DNA to 6.25 µL of 2X GoTaq Green Master Mix (Promega), 4.75 µL of nuclease-free water and 400 nM of each primer or by adding 1 µL of yeast lysate to 5 µL of DreamTaq Green PCR Master Mix (2X) (ThermoFisher Scientific) plus 4 µL of nuclease-free water with 200 nM of each primer. PCR reactions were performed as follows: 95°C for 2 min followed by 30 cycles of 95°C /2 min 30 s, 50°C /1 min 15 s, 72°C /1 min 45 s followed by a 72°C /5 min extension time (GoTaq) or 95°C for 2 min followed by 40 cycles of 95°C /20 s, 52°C /45 s, 72°C /1 min 30 s followed by a 72°C /2 min extension time (DreamTaq). 8-10 µL aliquots of each reaction were then run on a 1.5-2% TAE agarose gel.

### Promoter activity

Promoter activity was quantified as the ratio of YFP to mCherry fluorescence. Relative activity of native promoters was normalised to the standardised promoter *CYC1p* and OD₆₀₀.

### Ploidy determination by flow cytometry

Yeast ploidy was determined by measuring DNA content of ethanol-fixed cells stained with SYTOX Green and analyzed on a Sony SH800 cell sorter (Sony Biotechnology). Cells were grown to mid-log phase (OD₆₀₀ ≈ 0.5), and 2 mL of culture was harvested by centrifugation (2,000 × g, 4 min). Pellets were washed once with sterile filtered H₂O and fixed in 1 mL of 70% ethanol. Fixed cells were stored at 4 °C for at least 1 h. Cells were pelleted (2,000 × g, 4 min) and washed twice with 1 mL sterile 50 mM sodium citrate buffer. Samples were resuspended in 1 mL 50 mM sodium citrate containing RNase A (0.25 mg/mL final concentration) and incubated at 50 °C for 1 h. Proteinase K was then added to a final concentration of 0.4 mg/mL, and incubation continued for an additional 1 h at 50 °C. Cells were harvested again by centrifugation and resuspended in 1 mL 50 mM sodium citrate containing SYTOX Green (1:5,000 dilution from a 5 mM stock). Samples were incubated for at least 15 min at room temperature in the dark prior to analysis.Flow cytometry was performed using 488 nm excitation and FITC emission detection. A minimum of 10,000 events per sample was recorded. Forward and side scatter gating was applied to exclude debris and cell aggregates. Haploid (BY4742) and diploid (BY4743) control strains were included in each experiment as ploidy standards [76].s

### EASY-C procedure for mini-synthetic chromosome and plasmid

The EASY-C procedure for validation of mini-synthetic chromosomes and plasmids was performed as previously described (Swidah et al., 2026). Briefly, high-molecular-weight DNA was prepared and processed following the EASY-C workflow to enable structural verification and connectivity analysis of synthetic constructs. Library preparation, sequencing, and downstream analysis were conducted according to the published protocol without modification.

### High-molecular-weight DNA extraction

High-molecular-weight DNA was extracted using the NucleoBond HMW kit (Macherey-Nagel). Library preparation used the SQK-LSK109 kit (Oxford Nanopore Technologies). Sequencing was performed on a MinION Mk1B with Flongle flow cells. Data analysis used Guppy, NanoPlot, Minimap2, Sniffles, and Canu. Whole-plasmid sequencing was outsourced to Plasmidsaurus (UK).

### Long-read nanopore sequencing

Nanopore sequencing was performed to detect chromosome rearrangements (CRs), single nucleotide polymorphisms (SNPs), and insertion/deletion events in synthetic chromosomes. DNA purity and integrity were assessed prior to sequencing using agarose gel electrophoresis, a NanoDrop™ 2000 spectrophotometer, and a Qubit 4 fluorometer with dsDNA BR reagents (Thermo Fisher Scientific, USA) to ensure high-quality high-molecular-weight DNA. Circular mini-chromosomes were linearized using a restriction enzyme that cuts once within the plasmid backbone, and the digested DNA was purified using the QIAquick PCR Purification Kit (Qiagen, Germany). Libraries were prepared using the SQK-LSK109 Ligation Sequencing Kit together with Native Barcoding Kits EXP-NBD104 or EXP-NBD114 (Oxford Nanopore Technologies, UK). The protocol was modified by increasing the starting DNA input to approximately 2 µg without mechanical shearing to preserve long-read integrity. Sequencing was performed on a MinION Mk1B device equipped with a Flongle R9.4.1 flow cell (FLO-FLG001) for 24 h, targeting >20× genome coverage. Raw signal basecalling was conducted using Guppy v5.0.11 (Oxford Nanopore Technologies). Read quality was evaluated with NanoPlot v1.35.5. Reads were aligned to the reference genome using Minimap2 v2.20 or NGM-LR v0.2.7. Structural variants and indels were detected using Sniffles v1.0.12, and de novo assembly was performed with Canu v2.1.1. Whole-plasmid nanopore sequencing of selected constructs was additionally outsourced to Plasmidsaurus (UK).

## Supporting information

Figure 1 S. Post-SCRaMbLE analysis of first-generation SynIII + pRS-eNeochrome III.V1 S.c (1g) reveals few rearrangements

## Swidah et al figure captions

**Figure 1 S.**
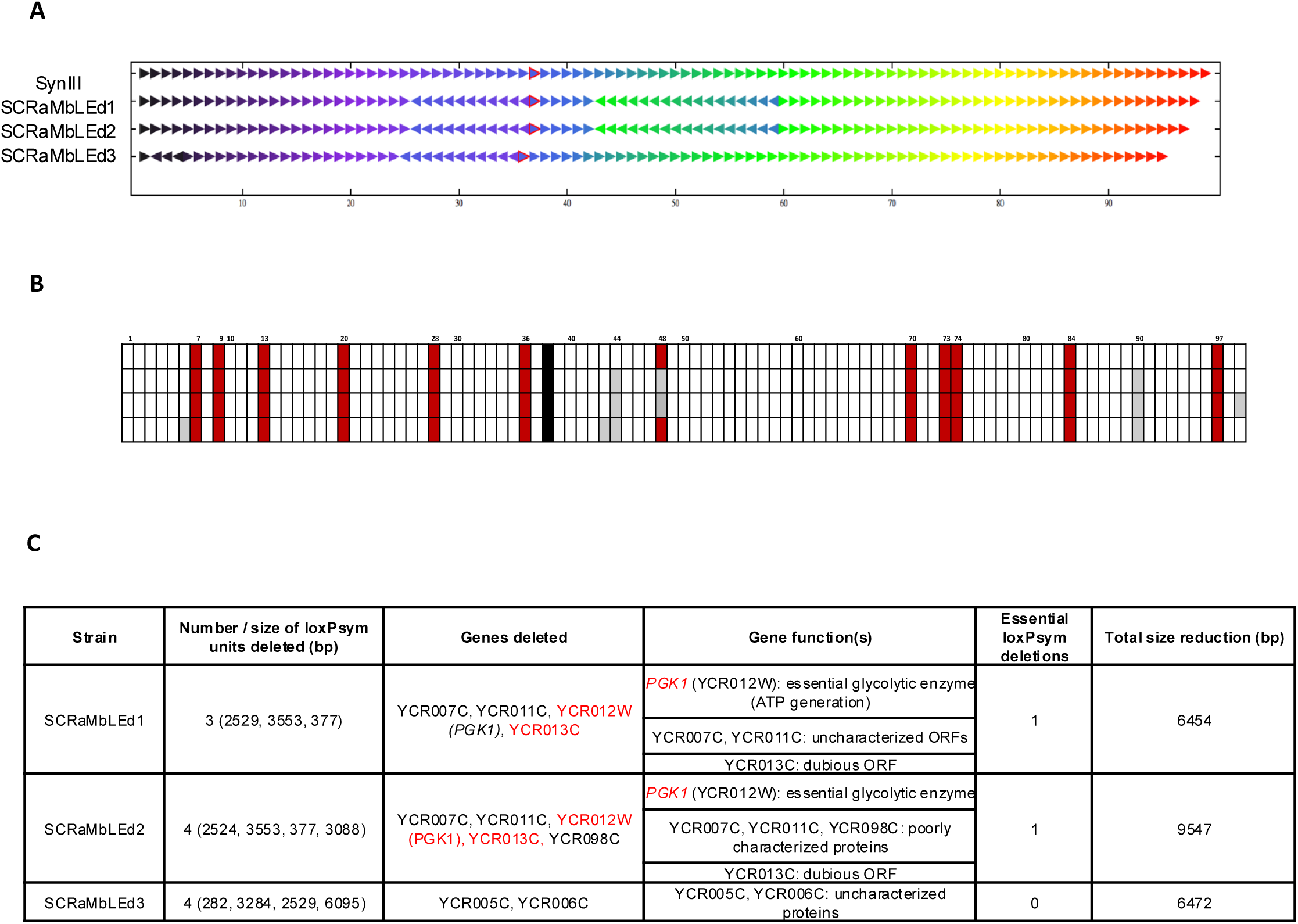
Post-SCRaMbLE analysis of first-generation SynIII + pRS-eNeochrome III.V1 S.c (1g) reveals few rearrangements. **A.** Arrow plot showing limited SCRaMbLEd rearrangements in SynIII + pRS-eNeochrome III.V1 S.c (1g) after 24 h induction of Cre recombinase under the control of the daughter-specific promoter pSCW11. The number of deletions increases from top to bottom. SynIII was used as the control strain. The centromere is represented by a red triangle, and each gene is represented by a triangle arranged from left to right. **B.** Schematic showing specific deletions in loxPsym units. SynIII serves as the control strain, with the centromere represented by a black box. Each loxPsym unit is shown as a box: essential regions are indicated in red and nonessential regions in white. The URA3 SCRaMbLE reporter is integrated at loxPsym site 44; all SCRaMbLEd strains lack the URA3 marker. SCRaMbLEd strains 1 and 2 exhibit deletion of essential loxPsym units, including essential genes such as PGK1, while YCR013C is a “dubious” open reading frame (∼7 kb). **C.** Summary table of key SCRaMbLE-induced deletions in post-SCRaMbLEd mutants of SynIII + pRS-eNeochrome III.V1 S.c (1g).

**Figure S2.**
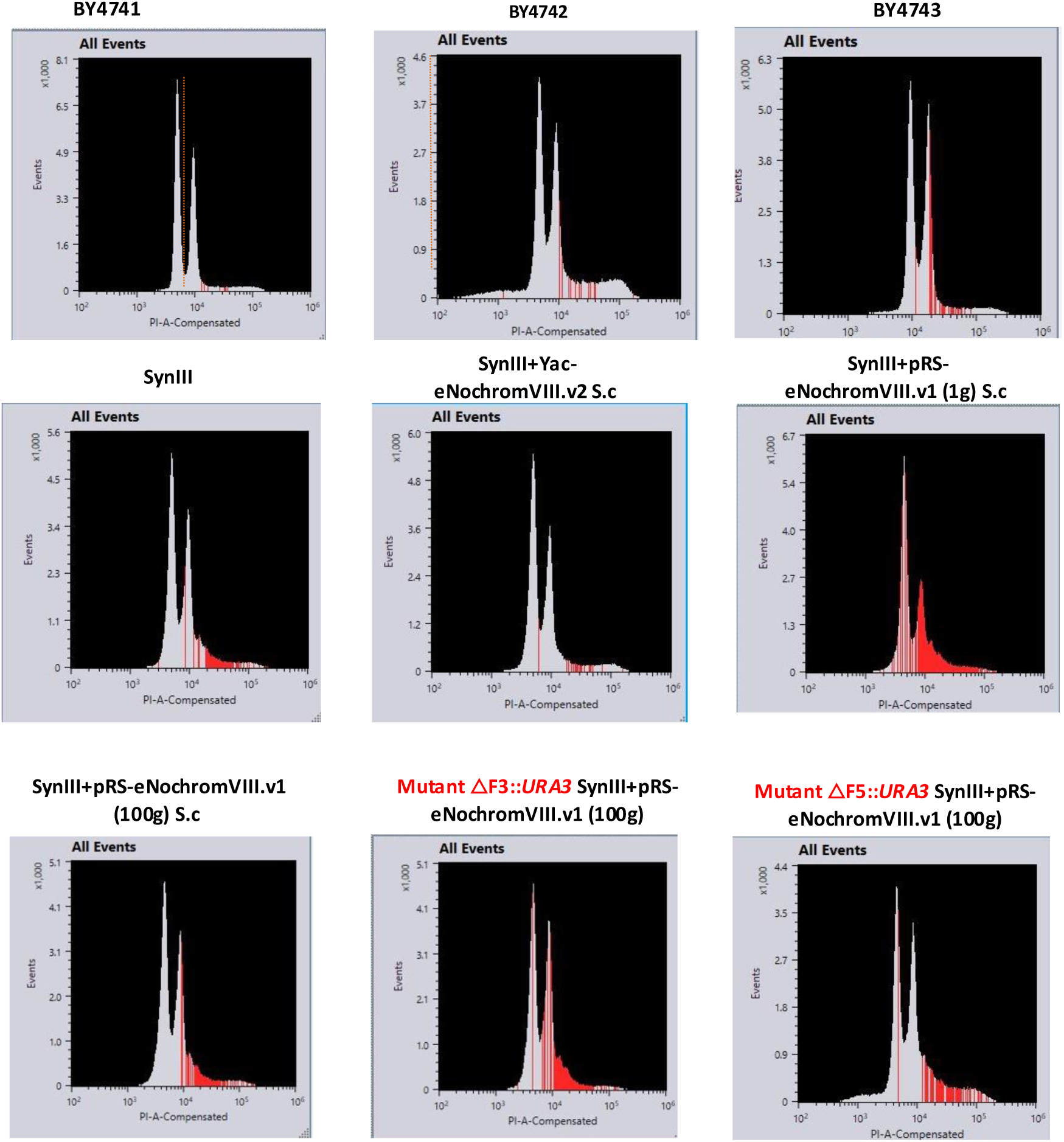
Evaluation of aneuploidy by FACS analysis. FACS analysis shows that synIII strains harboring different versions of eNeochrome, including potential mutants carrying deletion cassettes at various loci, are haploid. BY4741 (1n), BY4742 (1n), and BY4743 (2n) were used as control strains.

## Author contributions

RS (Investigation, Methodology, Project administration, Validation, Writing—original draft, Writing—review & editing, contributed to Funding acquisition); MM (Formal analysis of nanopore sequencing, Methodology, Validation, Writing—review & editing).

## Acknowledgments

We thank Professor Christopher Hardacre for support, including a strategic grant from the School of Natural Sciences at the University of Manchester. We acknowledge Dr. Eva Garcia-Ruiz for training in yeastFAB techniques, high-throughput workflows, and use of Echo and related liquid-handling systems, and for critical reading of the manuscript. We thank Dr. Stefan Hoffmann for providing wild-type mutants with orthogonal translation and Dr. Daniel Schindler for discussions at an early stage of the study.

## References

1. Kobayashi, K., et al., Essential Bacillus subtilis genes. Proc Natl Acad Sci U S A, 2003. 100(8): p. 4678–83.

2. Pena-Castillo, L. and T.R. Hughes, Why are there still over 1000 uncharacterized yeast genes? Genetics, 2007. 176(1): p. 7–14.

3. Serres, M.H., et al., A functional update of the Escherichia coli K-12 genome. Genome Biol, 2001. 2(9): p. RESEARCH0035.

4. Keseler, I.M., et al., The EcoCyc Database in 2021. Front Microbiol, 2021. 12: p. 711077.

5. Boone, C., H. Bussey, and B.J. Andrews, Exploring genetic interactions and networks with yeast. Nat Rev Genet, 2007. 8(6): p. 437–49.

6. Xu, X., et al., Trimming the genomic fat: minimising and re-functionalising genomes using synthetic biology. Nat Commun, 2023. 14(1): p. 1984.

7. Cello, J., A.V. Paul, and E. Wimmer, Chemical synthesis of poliovirus cDNA: generation of infectious virus in the absence of natural template. Science, 2002. 297(5583): p. 1016–8.

8. Dormitzer, P.R., et al., Synthetic generation of influenza vaccine viruses for rapid response to pandemics. Sci Transl Med, 2013. 5(185): p. 185ra68.

9. Gibson, D.G., et al., Complete chemical synthesis, assembly, and cloning of a Mycoplasma genitalium genome. Science, 2008. 319(5867): p. 1215–20.

10. Gibson, D.G., et al., Creation of a bacterial cell controlled by a chemically synthesized genome. Science, 2010. 329(5987): p. 52–6.

11. Fredens, J., et al., Total synthesis of Escherichia coli with a recoded genome. Nature, 2019. 569(7757): p. 514–518.

12. Zhao, Y., et al., Debugging and consolidating multiple synthetic chromosomes reveals combinatorial genetic interactions. Cell, 2023. 186(24): p. 5220–5236 e16.

13. Annaluru, N., et al., Total synthesis of a functional designer eukaryotic chromosome. Science, 2014. 344(6179): p. 55–8.

14. Luo, Z., et al., Compacting a synthetic yeast chromosome arm. Genome Biol, 2021. 22(1): p. 5.

15. Richardson, S.M., et al., Design of a synthetic yeast genome. Science, 2017. 355(6329): p. 1040–1044.

16. Wang, J., et al., Ring synthetic chromosome V SCRaMbLE. Nat Commun, 2018. 9(1): p. 3783.

17. Shen, Y., et al., SCRaMbLE generates designed combinatorial stochastic diversity in synthetic chromosomes. Genome Res, 2016. 26(1): p. 36–49.

18. Shen, Y., et al., Dissecting aneuploidy phenotypes by constructing Sc2.0 chromosome VII and SCRaMbLEing synthetic disomic yeast. Cell Genom, 2023. 3(11): p. 100364.

19. Schindler, D., et al., Design, construction, and functional characterization of a tRNA neochromosome in yeast. Cell, 2023.

20. Zhang, Z. and Q. Ren, Why are essential genes essential? - The essentiality of Saccharomyces genes. Microb Cell, 2015. 2(8): p. 280–287.

21. Umezu, K., et al., Purification and properties of orotidine-5’-phosphate pyrophosphorylase and orotidine-5’-phosphate decarboxylase from baker’s yeast. J Biochem, 1971. 70(2): p. 249–62.

22. van den Berg, M.A., et al., The two acetyl-coenzyme A synthetases of Saccharomyces cerevisiae differ with respect to kinetic properties and transcriptional regulation. J Biol Chem, 1996. 271(46): p. 28953–9.

23. Van den Berg, M.A. and H.Y. Steensma, ACS2, a Saccharomyces cerevisiae gene encoding acetyl-coenzyme A synthetase, essential for growth on glucose. Eur J Biochem, 1995. 231(3): p. 704–13.

24. Lygerou, Z., et al., The yeast BDF1 gene encodes a transcription factor involved in the expression of a broad class of genes including snRNAs. Nucleic Acids Res, 1994. 22(24): p. 5332–40.

25. Hartwell, L., Genetics. Robust interactions. Science, 2004. 303(5659): p. 774–5.

26. Sawa, C., et al., Bromodomain factor 1 (Bdf1) is phosphorylated by protein kinase CK2. Mol Cell Biol, 2004. 24(11): p. 4734–42.

27. Koonin, E.V., How many genes can make a cell: the minimal-gene-set concept. Annu Rev Genomics Hum Genet, 2000. 1: p. 99–116.

28. Mushegian, A.R. and E.V. Koonin, A minimal gene set for cellular life derived by comparison of complete bacterial genomes. Proc Natl Acad Sci U S A, 1996. 93(19): p. 10268–73.

29. Szathmary, E., Life: in search of the simplest cell. Nature, 2005. 433(7025): p. 469–70.

30. Hutchison, C.A., et al., Global transposon mutagenesis and a minimal Mycoplasma genome. Science, 1999. 286(5447): p. 2165–9.

31. Judson, N. and J.J. Mekalanos, TnAraOut, a transposon-based approach to identify and characterize essential bacterial genes. Nat Biotechnol, 2000. 18(7): p. 740–5.

32. French, C.T., et al., Large-scale transposon mutagenesis of Mycoplasma pulmonis. Mol Microbiol, 2008. 69(1): p. 67–76.

33. Ji, Y., et al., Identification of critical staphylococcal genes using conditional phenotypes generated by antisense RNA. Science, 2001. 293(5538): p. 2266–9.

34. Herring, C.D., J.D. Glasner, and F.R. Blattner, Gene replacement without selection: regulated suppression of amber mutations in Escherichia coli. Gene, 2003. 311: p. 153–63.

35. Giaever, G., et al., Functional profiling of the Saccharomyces cerevisiae genome. Nature, 2002. 418(6896): p. 387–91.

36. Winzeler, E.A., et al., Functional characterization of the S. cerevisiae genome by gene deletion and parallel analysis. Science, 1999. 285(5429): p. 901–6.

37. Costanzo, M., et al., A global genetic interaction network maps a wiring diagram of cellular function. Science, 2016. 353(6306).

38. Kuzmin, E., et al., Systematic analysis of complex genetic interactions. Science, 2018. 360(6386).

39. Luo, Z., et al., Whole genome engineering by synthesis. Sci China Life Sci, 2018. 61(12): p. 1515–1527.

40. Dymond, J. and J. Boeke, The Saccharomyces cerevisiae SCRaMbLE system and genome minimization. Bioeng Bugs, 2012. 3(3): p. 168–71.

41. Liu, W., et al., Rapid pathway prototyping and engineering using in vitro and in vivo synthetic genome SCRaMbLE-in methods. Nature Communications, 2018. G.

42. Ong, J.Y., et al., SCRaMbLE: A Study of Its Robustness and Challenges through Enhancement of Hygromycin B Resistance in a Semi-Synthetic Yeast. Bioengineering (Basel), 2021. 8(3).

43. Peng, B., et al., Controlling heterologous gene expression in yeast cell factories on different carbon substrates and across the diauxic shift: a comparison of yeast promoter activities. Microb Cell Fact, 2015. 14: p. 91.

44. Peng, B., et al., An Expanded Heterologous GAL Promoter Collection for Diauxie-Inducible Expression in Saccharomyces cerevisiae. ACS Synth Biol, 2018. 7(2): p. 748–751.

45. Artieri, C.G. and H.B. Fraser, Evolution at two levels of gene expression in yeast. Genome Res, 2014. 24(3): p. 411–21.

46. Fay, J.C. and J.A. Benavides, Evidence for domesticated and wild populations of Saccharomyces cerevisiae. PLoS Genet, 2005. 1(1): p. 66–71.

47. Kellis, M., B.W. Birren, and E.S. Lander, Proof and evolutionary analysis of ancient genome duplication in the yeast Saccharomyces cerevisiae. Nature, 2004. 428(6983): p. 617–24.

48. Sampaio, J.P., Microbe Profile: Saccharomyces eubayanus, the missing link to lager beer yeasts. Microbiology (Reading), 2018. 164(9): p. 1069–1071.

49. Hum, Y.F. and S. Jinks-Robertson, Mismatch recognition and subsequent processing have distinct effects on mitotic recombination intermediates and outcomes in yeast. Nucleic Acids Res, 2019. 47(9): p. 4554–4568.

50. Guo, Y., et al., YeastFab: the design and construction of standard biological parts for metabolic engineering in Saccharomyces cerevisiae. Nucleic Acids Res, 2015. 43(13): p. e88.

51. Garcia-Ruiz, E., et al., YeastFab: High-Throughput Genetic Parts Construction, Measurement, and Pathway Engineering in Yeast. Methods Enzymol, 2018. 608: p. 277–306.

52. Montrocher, R., et al., Phylogenetic analysis of the Saccharomyces cerevisiae group based on polymorphisms of rDNA spacer sequences. Int J Syst Bacteriol, 1998. 48 Pt 1: p. 295–303.

53. Ben-Aroya, S., et al., Toward a comprehensive temperature-sensitive mutant repository of the essential genes of Saccharomyces cerevisiae. Mol Cell, 2008. 30(2): p. 248–58.

54. Mitchell, L.A., et al., qPCRTag Analysis - A High Throughput, Real Time PCR Assay for Sc2.0 Genotyping. Jove-Journal of Visualized Experiments, 2015(99).

55. Swidah, R., M. Monti, and D. Delneri, EASY-C: Extraction and Analysis of Small Yeast Chromosomes-A rapid and universal platform for recovering artificial mini-chromosomes from synthetic Sc2.0 yeast and large plasmids from Saccharomyces cerevisiae and nonconventional yeast species. Synth Biol (Oxf), 2026. 11(1): p. ysag002.

56. Anastassiadis, K., et al., Dre recombinase, like Cre, is a highly efficient site-specific recombinase in E. coli, mammalian cells and mice. Dis Model Mech, 2009. 2(9-10): p. 508–15.

57. Murray, A.W. and J.W. Szostak, Construction of artificial chromosomes in yeast. Nature, 1983. 305(5931): p. 189–93.

58. Izvolsky, K.I., et al., Yeast artificial chromosome segregation from host chromosomes with similar lengths. Nucleic Acids Res, 1998. 26(21): p. 5011–2.

59. Guerrini, A.M., et al., Cloning a fragment from the telomere of the long arm of human chromosome 9 in a YAC vector. Chromosoma, 1990. GG(2): p. 138–42.

60. Pommier, Y., Topoisomerase I inhibitors: camptothecins and beyond. Nat Rev Cancer, 2006. 6(10): p. 789–802.

61. Wang, X.H., et al., Design, synthesis, and biological activity evaluation of campthothecin-HAA-Norcantharidin conjugates as antitumor agents in vitro. Chem Biol Drug Des, 2019. 93(6): p. 986–992.

62. Lundin, C., et al., Methyl methanesulfonate (MMS) produces heat-labile DNA damage but no detectable in vivo DNA double-strand breaks. Nucleic Acids Res, 2005. 33(12): p. 3799–811.

63. Luo, Z., S. Jiang, and J. Dai, Chromosomal Rearrangements of Synthetic Yeast by SCRaMbLE. Methods Mol Biol, 2021. 2196: p. 153–165.

64. Cherry, J.M., et al., Saccharomyces Genome Database: the genomics resource of budding yeast. Nucleic Acids Res, 2012. 40(Database issue): p. D700–5.

65. Goffeau, A., et al., Life with 6000 genes. Science, 1996. 274(5287): p. 546, 563–7.

66. Zhao, Y., et al., CREEPY: CRISPR-mediated editing of synthetic episomes in yeast. Nucleic Acids Res, 2023. 51(13): p. e72.

67. DiCarlo, J.E., et al., Genome engineering in Saccharomyces cerevisiae using CRISPR-Cas systems. Nucleic Acids Res, 2013. 41(7): p. 4336–43.

68. Chen, X.J. and G.D. Clark-Walker, The petite mutation in yeasts: 50 years on. Int Rev Cytol, 2000. 194: p. 197–238.

69. Costanzo, M., et al., The genetic landscape of a cell. Science, 2010. 327(5964): p. 425–31.

70. Wang, P., et al., SCRaMbLEing of a Synthetic Yeast Chromosome with Clustered Essential Genes Reveals Synthetic Lethal Interactions. ACS Synth Biol, 2020. 9(5): p. 1181–1189.

71. Gottschling, D.E., et al., Position effect at S. cerevisiae telomeres: reversible repression of Pol II transcription. Cell, 1990. 63(4): p. 751–62.

72. Guo, F., D.N. Gopaul, and G.D. van Duyne, Structure of Cre recombinase complexed with DNA in a site-specific recombination synapse. Nature, 1997. 389(6646): p. 40–6.

73. Dymond, J.S., et al., Synthetic chromosome arms function in yeast and generate phenotypic diversity by design. Nature, 2011. 477(7365): p. 471–6.

74. Gietz, R.D. and R.A. Woods, Transformation of yeast by lithium acetate/single-stranded carrier DNA/polyethylene glycol method. Methods Enzymol, 2002. 350: p. 87–96.

75. Sambrook, J. and D.W. Russell, Purification of nucleic acids by extraction with phenol:chloroform. CSH Protoc, 2006. 2006(1).

76. Haase, S.B. and S.I. Reed, Improved flow cytometric analysis of the budding yeast cell cycle. Cell Cycle, 2002. 1(2): p. 132–6.

