## Supplementary material for "Functional characterisation of an essential neo-chromosome III in Sc2.0 strain reveals opportunities and challenges for genome minimisation in Sc3.0": Figure 1 S. Post-SCRaMbLE analysis of first-generation SynIII + pRS-eNeochrome III.V1 S.c (1g) reveals few rearrangements

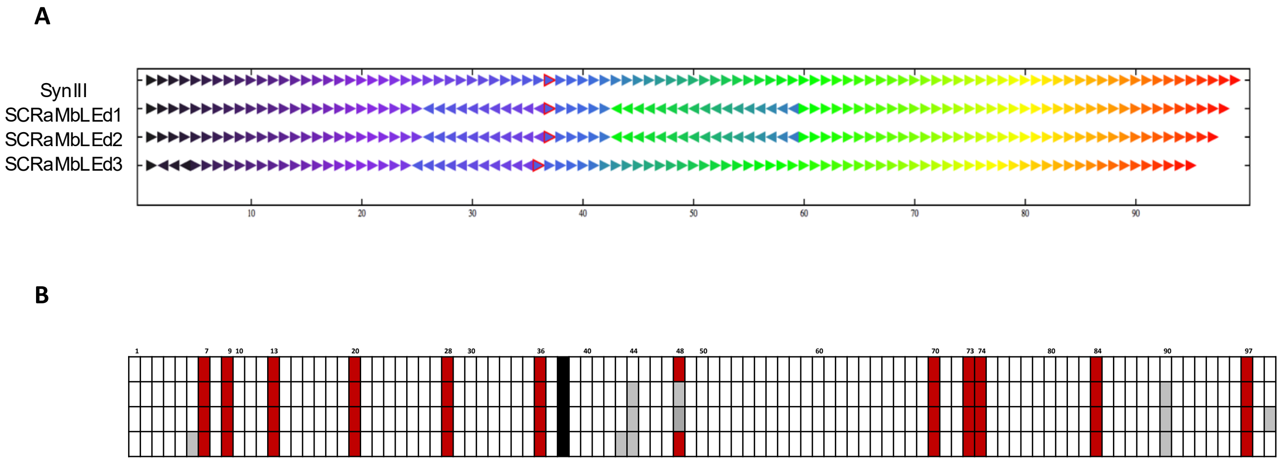


C

| Strain | Number / size of loxPsym units deleted (bp) | Genes deleted | Gene function(s) | Essential loxPsym deletions | Total size reduction (bp) |
|---|---|---|---|---|---|
| SCRaMbLEd1 | 3 (2529, 3553, 377) | YCR007C, YCR011C, YCR012W *(PGK1)*, YCR013C | *PGK1* (YCR012W): essential glycolytic enzyme (ATP generation)<br>YCR007C, YCR011C: uncharacterized ORFs<br>YCR013C: dubious ORF | 1 | 6454 |
| SCRaMbLEd2 | 4 (2524, 3553, 377, 3088) | YCR007C, YCR011C, YCR012W (PGK1), YCR013C, YCR098C | *PGK1* (YCR012W): essential glycolytic enzyme<br>YCR007C, YCR011C, YCR098C: poorly characterized proteins<br>YCR013C: dubious ORF | 1 | 9547 |
| SCRaMbLEd3 | 4 (282, 3284, 2529, 6095) | YCR005C, YCR006C | YCR005C, YCR006C: uncharacterized proteins | 0 | 6472 |

**Figure 2 S.** Evaluating aneuploidy using FACS analysis

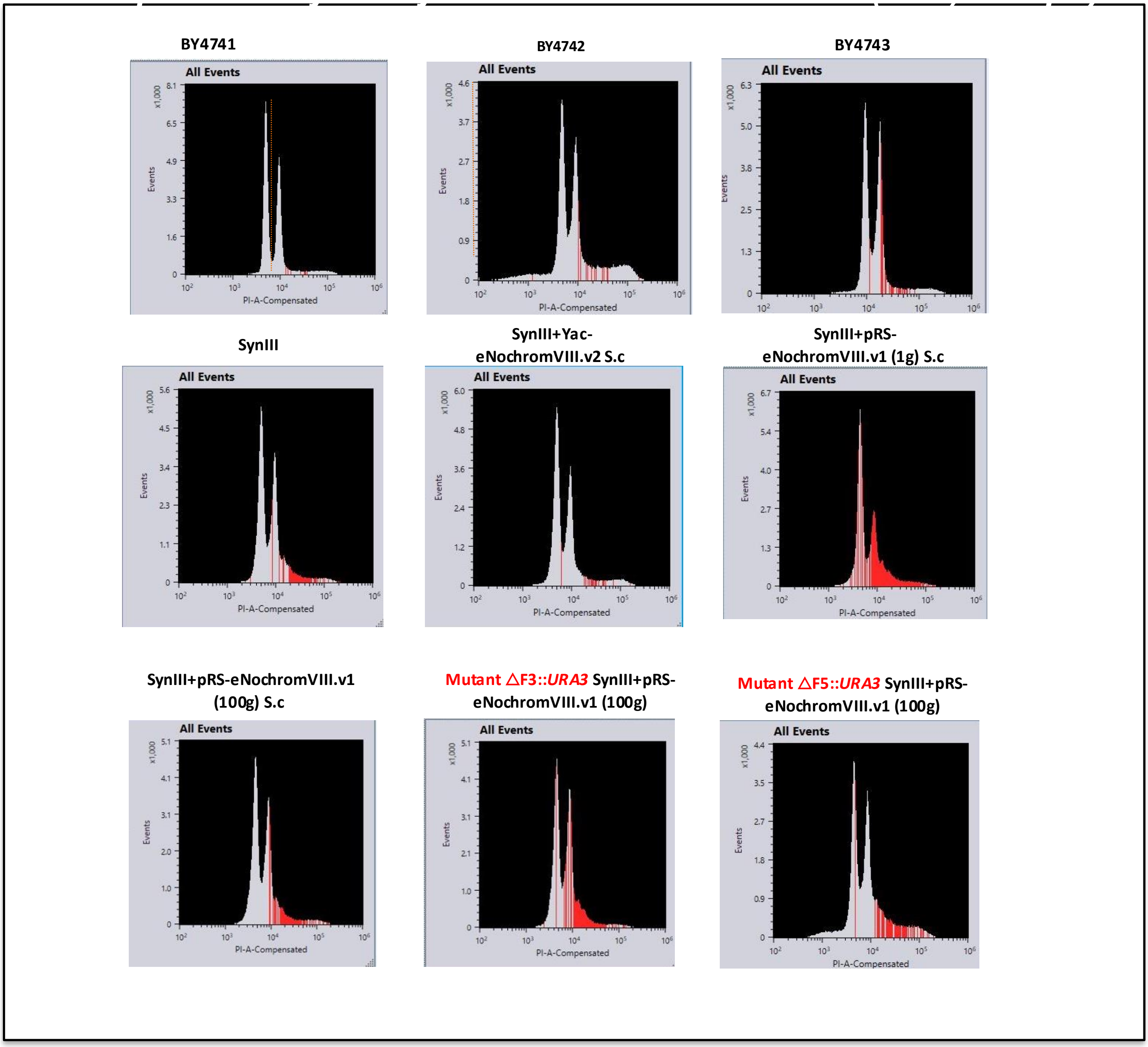
